# The tongue-brain axis mediates a hidden amino acid appetite

**DOI:** 10.64898/2026.04.17.719133

**Authors:** Shangming Wu, Yeting Gan, Min Tang, Shanghai Chen, Peixiang Luo, Kexin Tong, Kan Liu, Haizhou Jiang, Xiaoxue Jiang, Fei Xiao, Wei Lv, Feixiang Yuan, Feifan Guo

## Abstract

Selecting a diet containing all essential amino acids (EAAs) is critical for health. Following EAA deprivation, animals can select a nutritiously complete food source; however, the underlying mechanisms in vertebrates remain unclear. In mice, we show that leucine deficiency activates hypothalamic agouti-related protein (AgRP) neurons, which project to the paraventricular thalamus (PVT) via γ-aminobutyric acid and are required for EAA deficiency–induced leucine appetite in mice. Furthermore, the peripheral tongue amino acid sensor general control nonderepressive-2 (GCN2) mediates acute EAA appetite via AgRP neurons. Together, these findings identify a tongue–AgRP–PVT circuit underlying EAA appetite, which is important for the rapid and accurate selection of essential nutrients.

## Introduction

Diet is crucial for the survival of living animals. Animals have to decide the amount of food intake and select foods with different nutrients according to the internal or hedonic needs of the body ^1,2^. Carbohydrates, fats and proteins are the three major essential nutrients. Under normal circumstances, the body naturally has a preference for sugar and fat ^3^. This is partly attributable to the delicious food-based activation of the reward system to produce a sense of hedonism as well as to the body’s intrinsic need for nutrients ^2,3^. Proteins, which are composed of amino acids, are crucial for animal growth and development. However, the mechanisms underlying protein and amino acid appetite remain unclear. Under normal conditions, animals do not show a clear short-term preference for proteins or amino acids. However, under specific conditions, animals can develop a protein or amino acid appetite ^4-6^, including mated female fruit flies and protein- or amino acid–limited humans and rodents ^1,5-8^. Research advances on the mechanisms of protein/amino acid appetite in invertebrates have been reported primarily in fruit flies, such as the detection of the internal amino acid status by gustatory receptor neurons ^8,9^ and regulation of essential amino acid (EAA) appetite by gut general control nonderepressive-2 (GCN2) signaling via the microbiome–gut–brain axis ^1^. However, their regulatory mechanisms in vertebrates remain unelucidated, which limits therapeutic applications. In addition, it only takes a short time to decide what to eat for a meal or select preferable food ^10,11^. The ability to quickly decide what to eat is of great significance for survival, as the safe feeding time for animals in nature is limited and consuming inappropriate food may not meet the body’s needs. Whether animals can quickly detect and choose foods with needed amino acids, and the inside mechanisms have not been reported.

Agouti-related peptide (AgRP) neurons are crucial for sensing nutrients and integrating metabolic cues to regulate food intake ^12^. Under certain conditions, these neurons also mediate sugar and fat preference ^13-17^. Although AgRP neurons can sense leucine levels, their role in amino acid appetite and preference remains unknown ^18,19^. Leucine deprivation (and other EAA deprivations) can activate AgRP neurons, albeit with reduced food intake, which contradicts the orexigenic roles and implies unknown roles of AgRP neurons ^20,21^. Hence, we theorized that AgRP neurons were activated because of amino acid hunger (desiring the complete amino acid diets), which may mediate amino acid preference.

This study aimed to investigate whether leucine deprivation induces a strong amino acid appetite in mice, which can quickly recognize a complete diet, and whether the AgRP neurons with specific neural circuits mediate amino acid appetite.

## Results

### Leucine-deprivation induces a strong amino acid appetite in mice and enables rapid recognition of a complete diet

To systematically study whether mice that had been deprived of dietary amino acids were able to select food that contained complete amino acids and to develop an amino acid appetite, male C57BL/6J wild-type (WT) mice were subjected to a control or leucine-deficient diet for a period of time (from 1 h to 7 days) and then given a choice between the control diet and leucine-deficient diet for 12 h. The amino acid appetite was evaluated by the preference index—a commonly used measure to reflect food preference between two types of food ^1,22^. This index was calculated based on food-weight changes of control and leucine-deficient pellets consumed by the mice. Deprived male mice preferred a complete control diet over a leucine-deficient diet after only 1 h of leucine deprivation and the preference remained stable with 3-7 days of leucine-deprivation (**Fig. 1a**). We then used the 3-day leucine-deprivation treatment ^20,23^ to conduct subsequent analyses. To clarify how quickly the leucine-deficient treated mice could recognize the complete diet, WT mice were subjected to a control or leucine-deficient diet for 3 days and were then offered a choice between the control diet and leucine-deficient diet. For a 24-h record, the mice distinguished the control diet within 10 min and maintained the amino acid appetite longer (**Fig. 1b**). Similar results were observed in female mice (**Extended Data Fig. 1a**). As a leucine-deprived diet reduces food intake ^20^, we conducted experiments to investigate whether energy restriction could induce amino acid appetite. Control diet–fed mice, which received the same amount of food as the mice fed a leucine-deficient diet (pair-fed group), showed no preference for the complete diet in the choice assays (**Fig. 1c**). As the foods that were provided had different colors, we wondered whether the fast choice was attributed to color memory. The mice were subjected to a control or leucine-deficient diet for 3 days and were then given a choice between the control diet and leucine-deficient diet with the same green color or the opposite color, relative to the previous diets. Nonetheless, the mice could choose the control diet (correct food with complete amino acids) with no color interference (**Extended Data Fig. 1b** and **1c**). To clear whether the preference was direct to leucine, a single amino acid, the mice were given a choice between liquids—water or leucine solution. As expected, leucine-deprived mice showed a preference for the leucine solution (**Extended Data Fig. 1d**). These results show that, without being aided by color memory, the leucine-deprived mice could quickly recognize and choose foods or liquids that contained leucine.

**Fig. 1.**
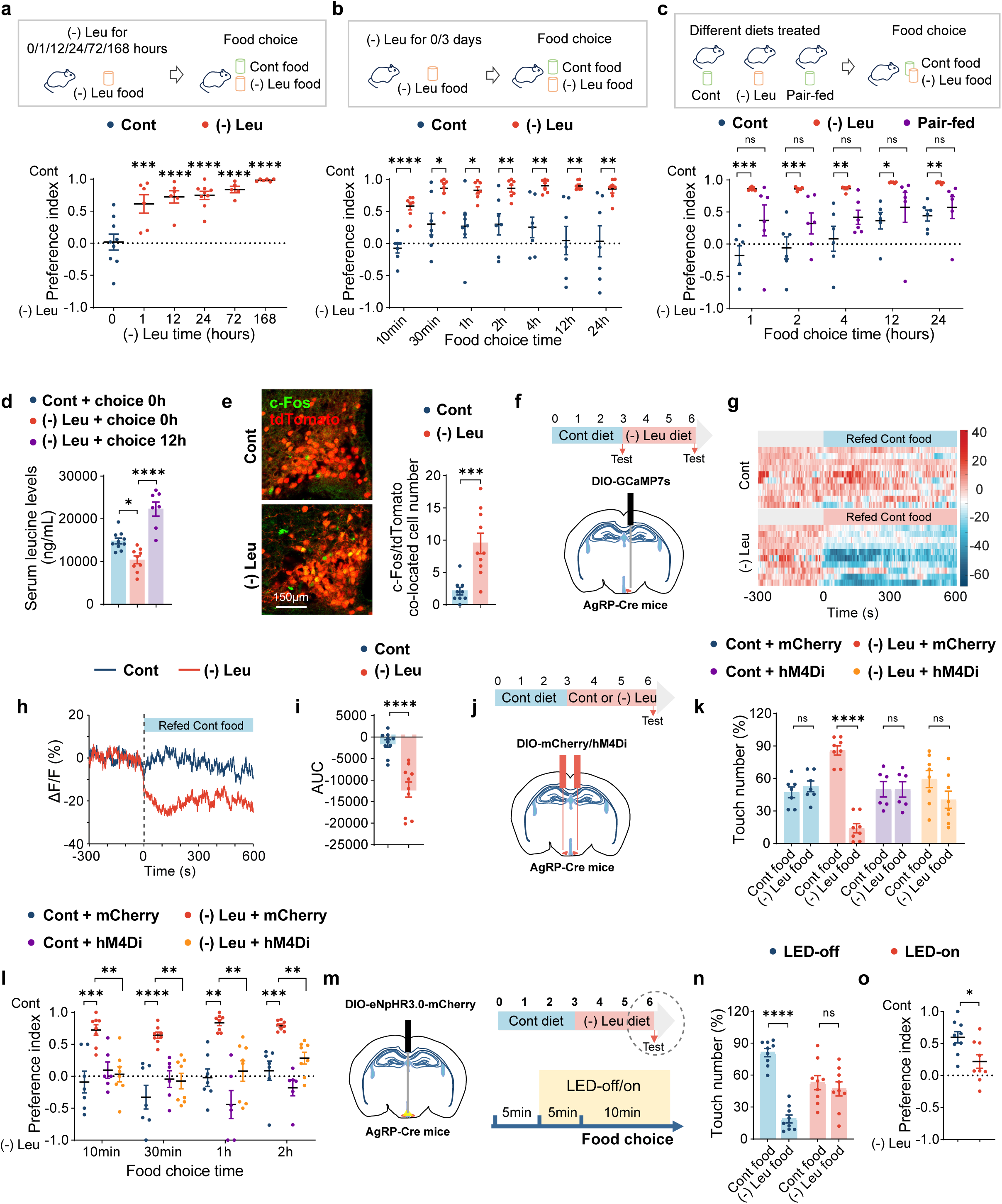
Hypothalamic AgRP neurons mediate leucine deprivation-induced amino acid appetite. a, Two-choice preferences for 12h of mice that fed a control (Cont) or leucine-deficient [(-) Leu] diet for indicated durations (1h, 12h, 24h, 72h and 168h). b, Two-choice preferences for indicated time (10min, 30min, 1h, 2h, 4h, 12h and 24h) of mice that fed a Cont or (-) Leu diet for 3 days. c,Two-choice preferences for indicated time (1h, 2h, 4h, 12h and 24h) of mice that fed a Cont, (-) Leu, or pair-fed diet for 3 days. d, Leucine concentration in serum of mice that fed a Cont or (-) Leu diet for 3 days and conducted two-choice preferences for 0 h or 12 h. e, Immunofluorescence (IF) staining for tdTomato (red), c-Fos (green) or merge (yellow) in arcuate nucleus (ARC) (left), and quantification of c-Fos and tdTomato colocalized cell numbers (right); 3V, third ventricle. f, Schematics illustrating the fiber photometry recordings of calcium signals in ARC^AgRP^ neurons and the timeline of diet feeding as well as the test. Each mouse was subjected to the experiment twice. g, Heatmap of the calcium signals in the ARC^AgRP^ neurons of mice that fed a Cont or (-) Leu diet for 3 days in response to a Cont diet. Each heatmap represents a single behavioral session. h, Averaged traces of calcium signals in g. i, The area under curve (AUC) of the calcium signals (0-600s) in h. j, Schematics illustrating virus-mediated mCherry/hM4Di expression (red) in ARC^AgRP^ neurons and the timeline of diet feeding as well as the test. Mice were fed a Cont or (-) Leu diet for 3 days followed by CNO injection. k, Touch number percentage for the first 10 min of two-choice assays in j. l, Two-choice preferences for indicated time (10min, 30min, 1h, 2h) in j. m, Schematics illustrating virus-mediated eNpHR expression (red) in ARC^AgRP^ neurons and the timeline of diet feeding as well as the photoinhibition test. Mice were fed a (-) Leu diet for 3 days, and the activity of AgRP was regulated by LED-off or LED-on. n, Touch number percentage for 10 min of two-choice assays in m. o, Two-choice preferences for 10min in m. Studies for a-d were conducted using 8–10-week-old male wild-type mice fed a Cont, (-) Leu, or pair-fed for 3 days or indicated durations; studies for e were conducted using 8–12-week-old male AgRP/Ai9 mice fed a Cont or (-) Leu diet for 3 days; studies for f-i were conducted using 12-16-week-old male AgRP-Cre mice receiving AAVs expressing GCaMP7s (AAV2/9-hSyn-DIO-jGCaMP7s) fed a Cont or (-) Leu diet for 3 days; studies for j - l were conducted using 12-16-week-old male AgRP-Cre mice receiving AAVs expressing mCherry or hM4Di fed a Cont or (-) Leu diet for 3 days; studies for m - o were conducted using 12-16-week-old female AgRP-Cre mice receiving AAVs expressing NpHR (AAV2/9-EF1-DIO-eNpHR3.0-mCherry) fed a (-) Leu diet for 3 days. Data are expressed as the mean ± SEM (n =5-10 per group, as indicated), with individual data points. Data were analyzed via one-way ANOVA with Dunnett’s multiple comparisons test (a, c), or one-way ANOVA with Tukey’s multiple comparisons test d, or two-tailed unpaired Student’s t-test (b, e, i, k, n, o), or two-way ANOVA followed by Tukey’s multiple comparisons test (l). *P < 0.05, **P < 0.01, ***P < 0.001, ****P < 0.0001.

Leucine deficiency reduces systemic leucine levels ^24^; therefore, we questioned whether the observed leucine preference was attributable to changes in leucine availability. We examined serum amino acids and found that leucine-deficient diets for 3 days resulted in low levels of leucine (**Fig. 1d**). However, following 12 h of food choice, leucine levels increased (**Fig. 1d**). This result shows that while decreased leucine may trigger leucine appetite and detection, it is not sufficient to sustain it. To assess whether the induced amino acid preference was leucine-specific, we conducted analogous experiments with other EAAs (tryptophan, valine, and isoleucine) and non-essential amino acids (NEAAs; glycine and glutamate). Mice fed different EAA-deficient diets, like those fed a leucine-deficient diet, preferred complete nutrient diets (**Extended Data Fig. 1e-1g**). However, mice fed single NEAA-deficient diets showed no preference for complete diets (**Extended Data Fig. 1h** and **1i**).

These results indicate that EAA deprivation leads to a fast and long-lasting EAA appetite, which is independent of sex, food ration, or diet color.

### Hypothalamic AgRP neurons mediate leucine deprivation–induced amino acid appetite

As hypothalamic AgRP neurons are crucial for appetite and nutrient preference, and can respond to leucine deprivation and leucine levels ^13,18,21^, we speculated that AgRP neurons participate in leucine deprivation–induced EAA appetite. We performed immunofluorescence (IF) staining to examine changes in c-Fos—a signal that reflects neuronal activity ^25^—in the AgRP neurons of AgRP-Ai9 reporter mice. IF staining revealed that c-Fos expression in AgRP neurons increased in leucine-deprivation-treated mice (**Fig. 1e**). Moreover, we investigated the dynamic regulation of AgRP neuronal activity in vivo using the Ca^2+^ sensor GCaMP7f in AgRP neurons. A Cre-dependent adeno-associated virus (AAV) encoding GCaMP7f (AAV2/9-DIO-GCaMP7f) was injected into the arcuate nucleus (ARC) of male AgRP-Cre mice, with an optical fiber implanted above the site of the AAVs (AgRP^GCaMP7f^; **Fig. 1f**). Four weeks after surgery, GCaMP7f signals were recorded. To measure the dynamic calcium signals of AgRP neurons in mice fed leucine-deficient and re-feeding control diets, AgRP^GCaMP7f^ mice were subjected to control or leucine-deficient diets for 3 days and were then transferred to control diets (**Fig. 1f**). We found that the AgRP neurons of mice treated with leucine-deficient diets exhibited a rapid and sustained decrease in calcium signals within seconds of being exposed to control diets. However, AgRP neurons of mice treated with control diets showed no significant signal changes (**Fig. 1g-1i**), which revealed that AgRP neurons were activated during the 3-day leucine deprivation.

Subsequently, we investigated the effect of chemogenetic inhibition of AgRP neural activity on leucine-deprivation-induced amino acid preference. Accordingly, we conducted inhibitory hM4Di designer receptors exclusively activated by designer drugs (DREADDs), which are activated by the inert ligand clozapine N-oxide (CNO). A Cre-dependent AAV encoding hM4Di (AAV2/9-DIO-hM4Di-mCherry) or mCherry (AAV2/9-DIO-mCherry) was injected into the ARC of male AgRP-Cre mice, which were then intraperitoneally (i.p.) injected with CNO before the experiments (**Fig. 1j**). Inhibition of AgRP neural activity was confirmed by reduced IF staining of c-Fos in AgRP neurons in mice injected with hM4Di (**Extended Data Fig. 2a**), in addition to reduced food intake after fasting, as AgRP neurons are required for the refeeding response (**Extended Data Fig. 2b**). The inhibition of AgRP neural activity largely blocked leucine-deprivation-induced leucine appetite, assessed by the percentage of the number of times the mice touched the different food, and the preference index (**Fig. 1k** and **1l**). Furthermore, to amplify the feeding effects, we conducted choice assays after 24 h of fasting. Mice pretreated with a leucine-deficient diet still showed an appetite for a leucine-containing diet, which could be blocked via the inhibition of AgRP neurons (**Extended Data Fig. 2c–2e**). To further verify the role of AgRP neurons in amino acid preference, we manipulated AgRP neural activity using an optogenetic approach by injecting AAV2/9-DIO-eNpHR-mCherry via optical fiber implantation (**Fig. 1m**). Yellow light (589nm) inhibited AgRP neural activity, as verified by the reduced food intake (**Extended Data Fig. 2f**). As expected, the optogenetic inhibition of AgRP neurons induced a reverse preference for complete amino acid foods within 10 min (**Fig. 1n** and **1o**).

To investigate whether the role of AgRP neurons in amino acid appetite is leucine specific, we inhibited AgRP neural activity using AAV-encoding hM4Di and conducted food-choice assays between the control and tryptophan-deficient diets. Mice that were fed tryptophan-deficient diets developed tryptophan appetite, which was blocked by inhibiting AgRP neural activity (**Extended Data Fig. 3**). These results show that the role of AgRP neurons in amino acid appetite is not specific to leucine.

In addition to AgRP neurons, pro-opiomelanocortin (POMC) neurons are important food-regulating neurons. We therefore examined their role in food choice and found that POMC neurons were activated during leucine deprivation (**Extended Data Fig. 4a**). To test their function, we inhibited POMC neural activity by injecting AAV2/9-DIO-hM4Di-mCherry into the ARC of POMC-Cre mice treated with CNO during the experiments (**Extended Data Fig. 4b–d**). However, inhibition of POMC neurons had no effect on leucine deprivation–induced amino acid preference (**Extended Data Fig. 4e** and **4f**).

These results indicate that hypothalamic AgRP neurons mediate leucine deprivation-induced amino acid appetite.

### Hypothalamic ARC^AgRP^-PVT^Glu^ neural circuits contribute to amino acid appetite

Subsequently, we investigated which AgRP terminals in specific brain regions promote amino acid appetite. AgRP neurons have widespread projections to different downstream sites in a one-to-one manner ^26^. Therefore, we used AgRP neuronal terminal-specific optogenetics to investigate the function of target areas without disturbing other branches. AgRP-Cre mice were injected with a Cre-dependent AAV encoding eNpHR (AAV2/9-DIO-eNpHR-mCherry) in ARC, and optic fibers were implanted into various brain regions that receive input from AgRP neurons, including the dorsomedial hypothalamic nucleus (DMH), bed nucleus of the stria terminalis (BNST), lateral hypothalamus (LH), paraventricular thalamus (PVT), dorsal raphe (DR), central amygdala (CeA), and hypothalamic paraventricular nucleus (PVN) (**Fig. 2a**). The efficiency of optogenetic inhibition of AgRP terminals was validated by the AAV and optic fiber sites (**Extended Data Fig. 5a**). Subsequently, the mice were subjected to leucine-deficient diets followed by fasting and food choice assays were conducted. Optogenetic inhibition of AgRP neuron terminals in the PVT, but not in other brain regions analyzed (DMH, BNST, LH, DR, CeA or PVN), blocked leucine deprivation–induced amino acid appetite (**Fig. 2b** and **2c**). Moreover, we used a Cre-dependent retro AAV encoding hM4Di (retroAAV-DIO-hM4Di-mCherry) injecting into the PVT of AgRP-Cre mice to confirm the projection from AgRP neuron to PVT (**Fig. 2d**). The results showed that inhibition of AgRP neurons projecting to PVT by CNO decreased c-Fos expression in AgRP neurons and attenuated leucine deprivation–induced amino acid appetite (**Fig. 2e**, **2f** and **Extended Data Fig. 5b**).

**Fig. 2.**
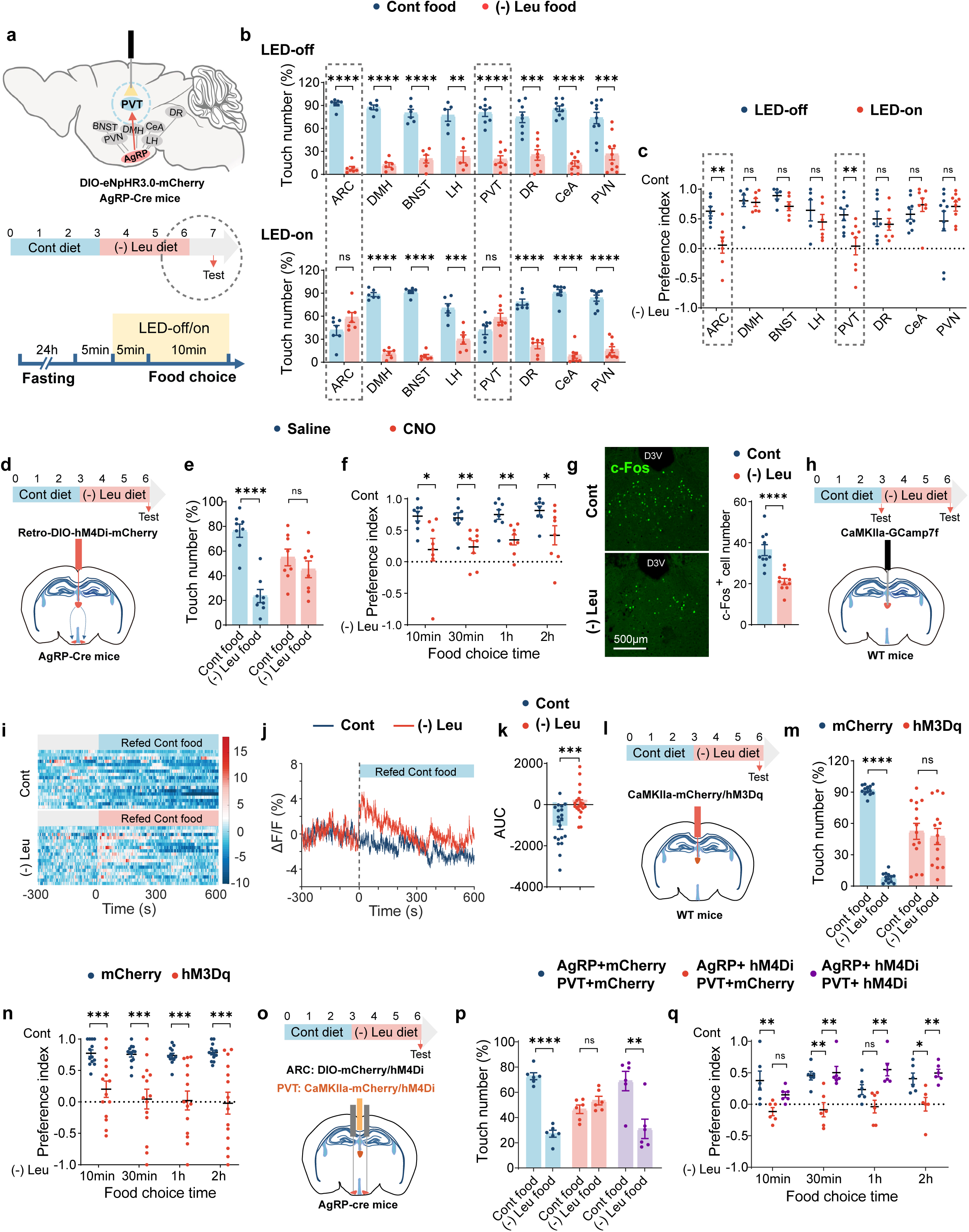
ARCAgRP-PVT^Glu^ neural circuit contributes to amino acid appetite. a, Schematics illustrating the locations of the optic fibre in ARC^AgRP^ neuron projections based on published brain section images (Top) and timeline of diet feeding, optogenetic stimulation process and two-choice preferences tests (Bottom). Mice were fed a leucine-deficient [(-) Leu] diet for 3 days, followed by LED-off or LED-on. b, Touch number percentage for 10 min of two-choice assays in (A), with LED-off (Top) and with LED-on (Bottom). c, Two-choice preferences for 10min in a. d, Schematics illustrating virus-mediated Retro-DIO-hM4Di-mCherry expression (red) in paraventricular thalamus (PVT) which traces to ARC^AgRP^ neurons and the timeline of diet feeding as well as the photoinhibition test. Mice were fed a (-) Leu diet for 3 days, and followed by saline or CNO injection. e, Touch number percentage for the first 10 min of two-choice assays in d. f, Two-choice preferences for indicated time (10min, 30min, 1h, 2h) in d. g, Immunofluorescence (IF) staining for c-Fos (green) in paraventricular thalamus (PVT) (left), and quantification of c-Fos cell numbers (right); D3V, dorsal third ventricle. h, Schematics illustrating the fiber photometry recordings of calcium signals in PVT^Glu^ neurons and the timeline of diet feeding as well as the test. Each mouse was subjected to the experiment twice. i, Heatmap of the calcium signals in the PVT^Glu^ neurons of mice that fed a Cont or (-) Leu diet for 3 days in response to a Cont diet. Each heatmap represents a single behavioral session. j, Averaged traces of calcium signals in (I). k, The area under curve (AUC) of the calcium signals (0-600s) in j. l, Schematics illustrating virus-mediated mCherry/hM3Dq expression (red) in PVT^Glu^ neurons and the timeline of diet feeding as well as the test. Mice were fed a (-) Leu diet for 3 days followed by CNO injection. m, Touch number percentage for the first 10 min of two-choice assays in l. n, Two-choice preferences for indicated time (10min, 30min, 1h, 2h) in l. o, Schematics illustrating virus-mediated mCherry/hM4Di expression (red) in PVT^Glu^ neurons, mCherry/hM4Di expression (red) in ARC^AgRP^ neurons, and the timeline of diet feeding as well as the test. Mice were fed a (-) Leu diet for 3 days followed by CNO injection. p, Touch number percentage for the first 10 min of two-choice assays in o. q, Two-choice preferences for indicated time (10min, 30min, 1h, 2h) in o. Studies for a-c were conducted using 8-14-week-old male AgRP-Cre mice [(dorsomedial hypothalamic nucleus (DMH), bed nucleus of the stria terminalis (BNST), PVT, and dorsa raphe (DR)], or using 12-16-week-old female AgRP-Cre mice [arcuate nucleus (ARC), lateral hypothalamus (LH), central amygdala (CeA), hypothalamic paraventricular nucleus (PVN)] receiving AAVs expressing NpHR (AAV2/9-EF1-DIO-eNpHR3.0-mCherry) with an optical fiber implantation into the indicated locations, fed a (-) Leu diet for 3 days; studies for d - f were conducted using 30-32-week-old male AgRP-Cre mice receiving AAVs expressing Retro-DIO-hM4Di-mCherry fed a (-) Leu diet for 3 days; studies for g were conducted using 8–12-week-old male wild-type (WT) mice fed a control or leucine-deficient diet for 3 days; studies for h-k were conducted using 10-12-week-old male WT mice receiving AAVs expressing GCaMP7f (AAV2/9-CaMKIIa-jGCaMP7f) in PVT fed a Cont or (-) Leu diet for 3 days; studies for l - n were conducted using 10-12-week-old male WT mice receiving AAVs expressing mCherry or hM3Dq in PVT fed a (-) Leu diet for 3 days; studies for o - q were conducted using 12-18-week-old male AgRP-Cre mice receiving AAVs expressing DIO-mCherry or DIO-hM4Di in ARC, and CaMKIIa-mCherry or CaMKIIa-hM4Di in PVT, fed a (-) Leu diet for 3 days. Data are expressed as the mean ± SEM (n = 5-20 per group, as indicated), with individual data points. Data were analyzed via two-tailed unpaired Student’s t-test (a-p), or one-way ANOVA followed by Tukey’s multiple comparisons test (q). *P < 0.05, **P < 0.01, ***P < 0.001, ****P < 0.0001.

Given the results that AgRP-PVT circuits contribute to amino acid appetite, we next investigated the role of PVT neurons in this context. IF staining revealed that leucine deprivation reduced the number of c-Fos in the PVT (**Fig. 2g**). The GCaMP7f signal (with CaMKIIα promoter, AAV2/9-CaMKIIα-GCaMP7f) recordings showed that the PVT glutamatergic (PVT^Glu^) neurons (as PVT neurons are predominantly glutamatergic) ^27^ exhibited a rapid increase in calcium signals after being switched from 3 d of leucine-deficient diet to the control diet (**Fig. 2h**-**2k**). To confirm whether PVT neurons contribute to the amino acid appetite, we injected the AAV-encoding hM3Dq with CaMKIIα promoter (AAV2/9-CaMKIIα-hM3Dq-mCherry) or mCherry (AAV2/9-CaMKIIα-mCherry) into the PVT of WT mice (**Fig. 2l**). The efficiency was validated using c-Fos staining and food intake after CNO injection (**Extended Data Fig. 6a** and **6b**), as previously reported ^28^. Stimulating PVT^Glu^ neurons abolished leucine deficiency–induced amino acid preference (**Fig. 2m** and **2n**). Furthermore, we questioned whether inhibition of PVT^Glu^ neurons could reverse AgRP inhibition-produced blocked effects on amino acid appetite. Therefore, the AAV2/9-DIO-hM4Di-mCherry or AAV2/9-DIO-mCherry was injected into the ARC, and the AAV2/9-CaMKIIα-hM4Di-mCherry or AAV2/9-CaMKIIα-mCherry was injected into the PVT of AgRP-Cre mice (**Fig. 2o**). The efficiency was confirmed by IF staining of c-Fos in AgRP and PVT neurons. Inhibition of AgRP neurons decreased c-Fos expression in AgRP neurons and increased it in PVT^Glu^ neurons, whereas inhibition of PVT^Glu^ neurons reduced c-Fos expression in PVT^Glu^ neurons (**Extended Data Fig. 6c**). Food-choice experiments showed that the suppression of AgRP neurons blocked leucine deficiency–induced amino acid preference, and this effect was reversed by PVT^Glu^ neuronal inhibition (**Fig. 2p** and **2q**). These results show that the AgRP-PVT^Glu^ circuits contributes to amino acid appetite.

### ARC^AgRP^ neurons act on PVT^Glu^ neurons via GABA signaling to modulate amino acid appetite

Next, we investigated how AgRP neurons acted on PVT^Glu^ neurons and whether any of the three principal neurotransmitters released by AgRP neurons—AgRP, NPY, or γ-aminobutyric acid (GABA) ^29^ —were required for leucine deprivation–induced amino acid preference. We examined the relative mRNA levels of these transmitters in ARC, including *Agrp*, *Npy*, *Gad1* (the key enzyme that generates GABA), and *Slc32a1* (also named *Vgat*, the transporter responsible for GABA transport) ^30^, under leucine deprivation and found that they were all increased (**Extended Data Fig. 7a**). We then tested the mRNA levels of AgRP, NPY, and GABA receptors in the PVT. GABA-B receptor 1, *Gabbr1* (the gene encoding GABAbR1), and the AgRP-acting receptor *Mc4r* increased in the leucine-deficient diet, whereas other receptors showed no change after the leucine-deficient diet (**Extended Data Fig. 7b**). To determine which neurotransmitter receptor mediated amino acid preference, we first injected two types of GABA-B receptor antagonists (Saclofen and CGP55845) into the PVT of WT mice (**Fig. 3a**). The injection of GABA-B receptor antagonists increased PVT c-Fos levels and blocked leucine deprivation–induced amino acid appetite (**Fig. 3b**, **3c** and **Extended Data Fig. 7c**). Furthermore, injections of a GABA-A receptor antagonist (bicuculline), AgRP receptor MC3/4R antagonist (SHU9119), and NPY 1/2/5R antagonist (BIBO3304, BIIE-0246, and CGP71683) into the PVT of WT mice had no influence on leucine deprivation–induced amino acid appetite (**Extended Data Fig. 7d–l**).

**Fig. 3.**
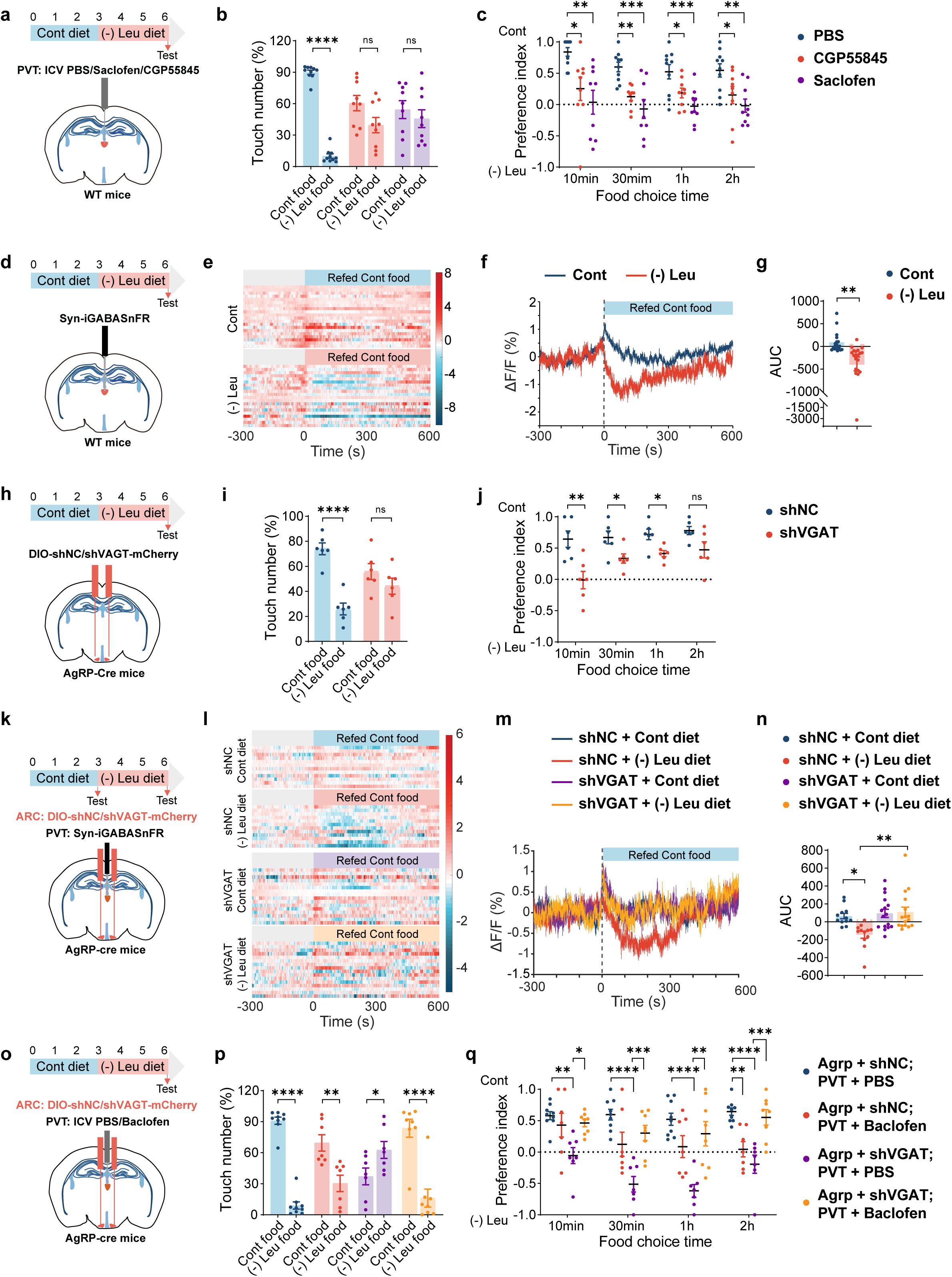
ARCAgRP neurons act on PVT^Glu^ neurons via GABA signaling to modulate amino acid appetite. a, Schematics illustrating PVT injection of Saclofen, CGP55845 or PBS, and the timeline of diet feeding as well as the test. Mice were fed a (-) Leu diet for 3 days followed by medicine injection. b, Touch number percentage for the first 10 min of two-choice assays in a. c, Two-choice preferences for indicated time (10min, 30min, 1h, 2h) in a. d, Schematics illustrating the fiber photometry recordings of GABA signals in PVT neurons and the timeline of diet feeding as well as the test. Each mouse was subjected to the experiment twice. e, Heatmap of the fluorescence signals in the PVT neurons of mice that fed a Cont or (-) Leu diet for 3 days in response to a Cont diet. Each heatmap represents a single behavioral session. f, Averaged traces of fluorescence signals in e. g, The area under curve (AUC) of the fluorescence signals (0-600s) in f. h, Schematics illustrating virus-mediated shVGAT expression (red) in ARC^AgRP^ neurons and the timeline of diet feeding as well as the test. Mice were fed a (-) Leu diet for 3 days. i, Touch number percentage for the first 10 min of two-choice assays in h. j, Two-choice preferences for indicated time (10min, 30min, 1h, 2h) in h. k, Schematics illustrating virus-mediated shVGAT expression (red) in ARC^AgRP^ neurons, the fiber photometry recordings of GABA signals in PVT neurons, and the timeline of diet feeding as well as the test. Each mouse was subjected to the experiment twice. l, Heatmap of the fluorescence signals in the PVT neurons of mice that fed a Cont or (-) Leu diet for 3 days in response to a Cont diet. Each heatmap represents a single behavioral session. m, Averaged traces of fluorescence signals in l. n, The area under curve (AUC) of the fluorescence signals (0-600s) in m. o, Schematics illustrating virus-mediated shVGAT expression (red) in ARC^AgRP^ neurons, the PVT injection of Baclofen or PBS, and the timeline of diet feeding as well as the test. Mice were fed a (-) Leu diet for 3 days. p, Touch number percentage for the first 10 min of two-choice assays in o. q, Two-choice preferences for indicated time (10min, 30min, 1h, 2h) in o. Studies for a-c were conducted using 10-12-week-old male WT mice with PVT injection of Saclofen, CGP55845 or PBS, fed a (-) Leu diet for 3 days; studies for d-g were conducted using 10-12-week-old male WT mice receiving AAVs expressing Syn-iGABASnFR fed a (-) Leu diet for 3 days; studies for h - j were conducted using 8-12-week-old female AgRP-Cre mice receiving AAVs expressing DIO-shNC or DIO-shVGAT in ARC fed a Cont or (-) Leu diet for 3 days; studies for k-n were conducted using 10-12-week-old male AgRP-Cre mice receiving AAVs expressing DIO-shNC or DIO-shVGAT in ARC, and Syn-iGABASnFR in PVT, fed a Cont or (-) Leu diet for 3 days; studies for o-q were conducted using 12-20-week-old female AgRP-Cre mice receiving AAVs expressing DIO-shNC or DIO-shVGAT in ARC, and PVT injection of Baclofen or PBS, fed a (-) Leu diet for 3 days. Data are expressed as the mean ± SEM (n = 6-20 per group, as indicated), with individual data points. Data were analyzed via two-tailed unpaired Student’s t-test (b, g, i, j, p), or one-way ANOVA followed by Dunnett’s multiple comparisons test (c), or two-way ANOVA followed by Tukey’s multiple comparisons test (n, q). *P < 0.05, **P < 0.01, ***P < 0.001, ****P < 0.0001.

Therefore, we focused on GABA, which may be a special neurotransmitter in AgRP neurons that regulates amino acid appetite ^31,32^. First, we injected a GABA probe (AAV2/9-syn-iGABASnFR-WPRE) into the PVT and performed fiber recordings (**Fig. 3d**). GABA signal recordings showed that PVT neurons exhibited a significant decrease in GABA signals after transfer to control diets following 3-day leucine-deficient diets, which showed that leucine deprivation increased the PVT neural extracellular GABA quantity, which was consistent with the increased activity of AgRP neurons (**Fig. 3e-g**). We then ascertained whether the PVT GABA signal changes were dependent on AgRP neurons. To inhibit GABA release in AgRP neurons, a Cre-dependent AAV encoding a short hairpin RNA directed against VGAT (AAV2/9-DIO-shVGAT-mCherry) or mCherry (AAV2/9DIO-shNC-mCherry) was injected into the ARC of AgRP-Cre mice (**Fig. 3h**). IF staining of mCherry (reflecting AgRP neurons) and VGAT revealed that VGAT was colocalized with AgRP neurons in control mice, but this co-localization was significantly reduced in AgRP-shVGAT mice (**Extended Data Fig. 7m**). The mRNA expression of *Vgat* in the ARC was lower in AgRP-shVGAT mice, as well as *Gad1*, with no changes in *Agrp* or *Npy* mRNA (**Extended Data Fig. 7n**). Knockdown of VGAT in AgRP neurons blocked leucine-deprivation-induced amino acid appetite (**Fig. 3i** and **3j**). In agreement with this, fiber recordings of the GABA probe showed that knocking down VGAT in AgRP neurons stopped the GABA signaling changes in PVT neurons after diet transfer from leucine-deficient diets (**Fig. 3k**-**3n**). In addition, knocking down AGRP or NPY in AgRP neurons had no obvious effect on amino acid preference (**Extended Data Fig. 7o-q**).

To further confirm the effects of AgRP secretory GABA and PVT GABA-B receptor, we injected AAV2/9-DIO-shVGAT-mCherry into the ARC in AgRP-Cre mice, and 4 weeks later, we injected a GABA-B receptor activator (Baclofen) into the PVT (**Fig. 3o**). Food-choice experiments showed that activating GABA-B receptors in the PVT reversed the effects of AgRP VGAT knockdown on amino acid preference (**Fig. 3p** and **3q**). These data reveal that the AgRP neurotransmitter GABA is required for amino acid preference, and that PVT GABA-B receptor mediates AgRP-PVT^Glu^ circuits in leucine deficiency-induced amino acid appetite.

### The tongue delivers nutrient signals to AgRP neurons to develop amino acid appetite

Subsequently, we questioned how food signals were transmitted to AgRP neurons to facilitate choice between full-nutrient diet and leucine-deprived diet. As the AgRP neurons can sense amino acid levels ^18^, we first assumed that hypothalamic AgRP neurons directly sense leucine signals to make the choice. However, injecting leucine into the third ventricle did not change the leucine deficiency–induced amino acid appetite (**Extended Data Fig. 8a–c**).

As mice developed an amino acid appetite and made the choice quickly within minutes after leucine-deficient treatment, we focused on the tongue—the organ that first comes into contact with food. As GCN2 is an amino acid sensor under EAA deprivation ^33^, we questioned whether GCN2 signaling plays a role in the tongue. As expected, GCN2 downstream signal p-eIF2a levels, amino acid-sensing gene *Asns* mRNA levels, and uncharged leucine tRNA levels increased in the tongue under leucine deprivation (**Fig. 4a**, **4b** and **Extended Data Fig. 8d**). We then knocked down GCN2 in the tongue by injecting AAVs encoding short-hairpin RNA directed against GCN2 (AAV2/DJ-shGCN2-EGFP) or GFP (AAV2/DJ-shNC-EGFP) into the tongue of AgRP-Ai9 mice (**Fig. 4c**, **4d** and **Extended Data Fig. 8e**). Surprisingly, tongue GCN2 knockdown reversed the amino acid preference within the first 10 min (**Fig. 4e** and **4f**). We then tested related signals in AgRP neurons. Knockdown of GCN2 in tongue inhibited AgRP neural activity under leucine deprivation, as validated by c-Fos staining and dynamic calcium signals of AgRP neurons (**Fig. 4g-k**), and reduced *Slc32a1* levels in the ARC (**Extended Data Fig. 8f**). Further, we knocked down GCN2 in the tongue and tested GABA signals in the PVT after leucine deprivation and refeeding with control diets (**Extended Data Fig. 8g**). These results showed that GCN2 knockdown in the tongue blocked GABA changes in PVT after leucine-deficient treatment (**Extended Data Fig. 8h-j**). These results show that tongue GCN2 is required for the recognition of leucine deprivation–induced amino acid preference and regulates AgRP neural activity.

**Fig. 4.**
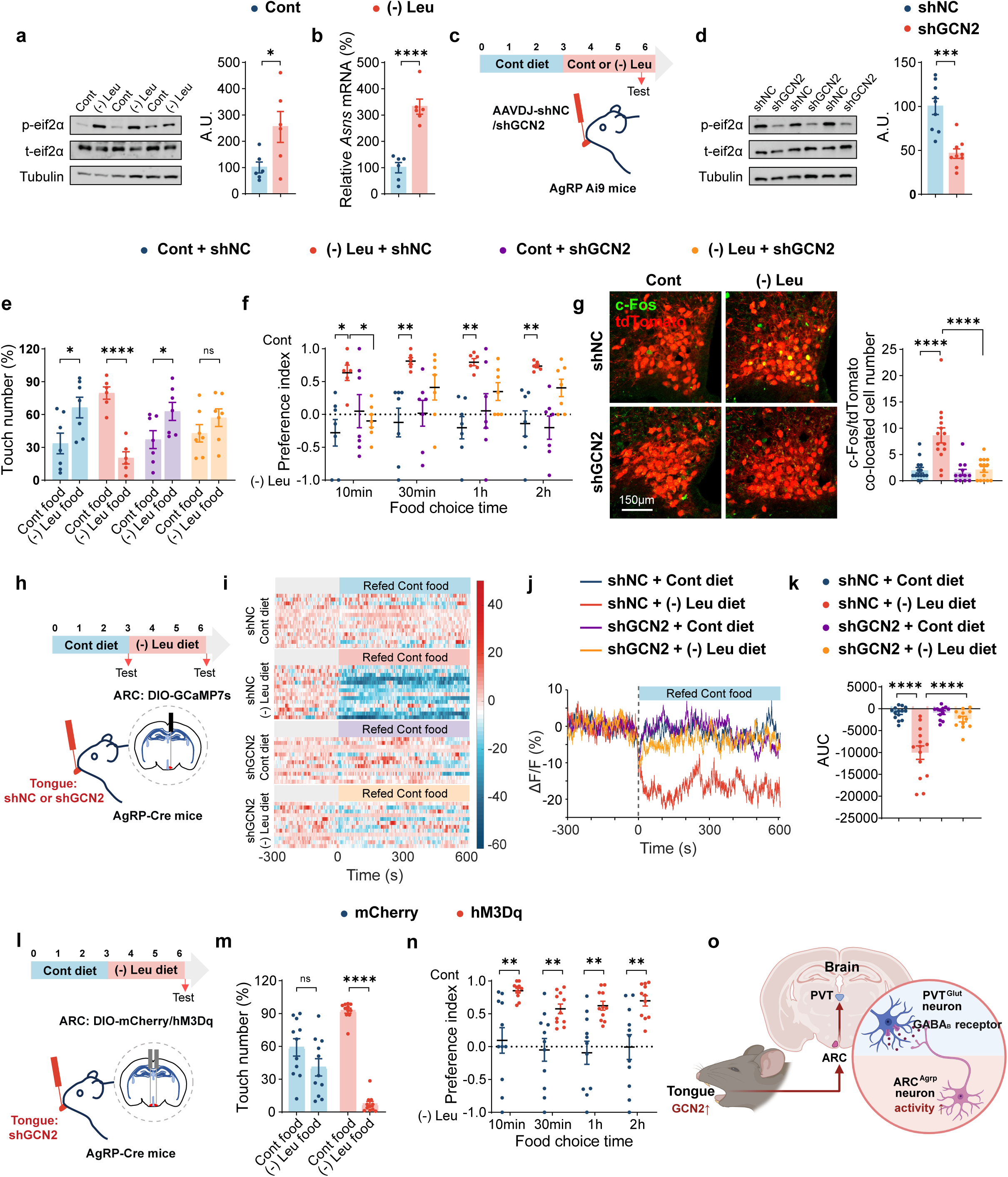
The tongue GCN2 signals act on AgRP neurons to develop amino acid appetite. a, p-eIF2α, t-eIF2α, and tubulin protein in the tongue by western blotting (left) and quantified by densitometric analysis (right), A.U.: arbitrary units. b, Gene expression of *Asns* in the tongue by RT-PCR. c, Schematics illustrating virus-mediated shGCN2-expression in the tongue of AgRP/Ai9 mice, and the timeline of diet feeding as well as the test. Mice were fed a Cont or (-) Leu diet for 3 days. d, p-eIF2α, t-eIF2α, and tubulin protein in the tongue by western blotting (left) and quantified by densitometric analysis (right). e, Touch number percentage for the first 10 min of two-choice assays in c. f, Two-choice preferences for indicated time (10min, 30min, 1h, 2h) in c. g, IF staining for tdTomato (red), c-Fos (green) or merge (yellow) in ARC (left), and quantification of c-Fos and tdTomato colocalized cell numbers (right). h, Schematics illustrating the fiber photometry recordings of calcium signals in ARC^AgRP^ neurons of AgRP-Cre mice receiving shGCN2 or shNC in the tongue and the timeline of diet feeding as well as the test. Each mouse was subjected to the experiment twice. i, Heatmap of the calcium signals in the PVT neurons of mice that receiving shGCN2 or shNC in the tongue fed a Cont or (-) Leu diet for 3 days in response to a Cont diet. Each heatmap represents a single behavioral session. j, Averaged traces of calcium signals in i. k, The area under curve (AUC) of the calcium signals (0-600s) in j. l, Schematics illustrating virus-mediated shGCN2 (red) in the tongue and DIO-mCherry or DIO-hM3Dq in the ARC, and the timeline of diet feeding as well as the test. m, Touch number percentage for the first 10 min of two-choice assays in l. n, Two-choice preferences for indicated time (10min, 30min, 1h, 2h) in l. o, Summary diagram: Short-duration EAA deficiency induces quick and continuous amino acid appetite, whereby quick amino acid detection is mediated by a tongue-AgRP-PVT neural circuit. Tongue GCN2 signaling is necessary for appetite and sends nerve signals to activate AgRP neurons. Created in BioRender. Shangming, W. (2026) https://BioRender.com/lxw97nn. Studies for a and b were conducted using 8-10-week-old male wild-type (WT) mice fed a Cont or (-) Leu diet for 3 days; studies for c-g were conducted using 40-48-week-old male AgRP/Ai9 mice receiving AAVs expressing shGCN2 or shNC in the tongue, fed a Cont or (-) Leu diet for 3 days; studies for h-k were conducted using 12-14-week-old male AgRP-Cre mice receiving AAVs expressing shGCN2 or shNC in the tongue and GCaMP7s (AAV2/9-hSyn-DIO-jGCaMP7s) in ARC, fed a Cont or (-) Leu diet for 3 days; studies for l-n were conducted using 12-15-week-old male AgRP-Cre mice receiving AAVs expressing shGCN2 in the tongue, and DIO-mCherry or DIO-hM4Di in ARC, fed a (-) Leu diet for 3 days. Data are expressed as the mean ± SEM (n = 6-14 per group, as indicated), with individual data points. Data were analyzed via two-tailed unpaired Student’s t-test (a, b, d, e, m, n), or two-way ANOVA followed by Tukey’s multiple comparisons test (f, g, k). *P < 0.05, **P < 0.01, ***P < 0.001, ****P < 0.0001.

Next, we suspected that the tongue had a neural connection with AgRP neurons. We conducted forward and retrograde tracing experiments to test this hypothesis. First, we injected forward tracing HSV (HSV-EGFP) into the tongue and retrograde tracing PRV (PRV-GAG-mCherry) into the ARC of WT mice (**Extended Data Fig. 9a**). We found that neurons in the ARC, ventromedial hypothalamic nucleus (VMH), and nucleus tractus solitarius (NTS; the brain region upstream of AgRP neurons from the periphery) ^34^ were traced by forward HSV, and tongue cells were traced by retrograde PRV (**Extended Data Fig. 9b**). Next, we injected forward-tracing HSV (HSV-EGFP) into the tongue and Cre-dependent retrograde-tracing PRV (PRV-DIO-GAG-mCherry) into the ARC of AgRP-Cre mice (**Extended Data Fig. 9c**). We found that AgRP neurons and the VMH were traced by forward HSV, and tongue cells were traced by retrograde PRV (**Extended Data Fig. 9d**).

To confirm the function of tongue-ARC^AgRP^ circuits, we injected AAV-shGCN2 into the tongue and AAV-DIO-hM3Dq into the AgRP neurons (**Fig. 4l**). The results showed that activation of the AgRP neurons can rescue the GCN2-knockdown-caused insensitivity in amino acid appetite (**Fig. 4m** and **4n**).

These results show that the tongue delivers nutrient signals to AgRP neurons to develop amino acid appetite.

## Discussion

Shifts in food preferences driven by nutrient-specific hunger are essential for survival, yet the underlying mechanisms in vertebrates remain poorly understood. Animal preferences for sugar and fat are direct and common ^2,35^. Amino acid preference seems special because amino acid or protein appetite is imperceptible and can be induced under certain conditions ^8,36^. Animals with EAA deprivation pre-treatment exhibited strong EAA appetite ^7,37^. For the mechanism, Animals can continuously detect sugars and glucose in various ways, including sensory experience, eating, ingestion and neurotransmission ^38-45^. Amino acid sensing may have similar mechanisms but specific research is limited. In this study, we further systematically investigated the factors related to the amino acid appetite and identified a tongue–brain neural circuit of hidden amino acid appetite which induced by EAA deficiency (**Fig. 4o**). The results showed that the induced amino acid appetite was EAA specific, quickly induced, quickly recognized, and long lasting, but not influenced by gender, color memory, food form, or serum amino acid levels. We then focused on early amino acid appetite, which is important for initiating meals but has received relatively little attention in previous research. To optimize fitness, animals must dynamically match food choices to their current needs ^9^. In our study and other studies, after EAA deprivation, the mice can quickly recognize the control food because of the reduced EAA levels in the body (serum), which is their internal needs ^1,9^. Our research has identified the peripheral-neural mechanism in animals that generates this intrinsic need and subsequently leads to the formation of preferences.

AgRP neurons are located in the hypothalamic ARC and have multiple functions, including feeding, food seeking, anti-depression, and learning ^46^. Among these, feeding (hunger) regulation appears to be the most common. Here, we found that the AgRP neuron regulates feeding and mediates “smart” feeding, rapidly detecting a complete amino acid diet and helping in making the right decision (choice). Our results expand on the neural function of AgRP through a detailed investigation of its role in amino acid sensing. We assumed that AgRP neurons sense diverse nutrients from multiple organs and play different roles through distributed circuits and special neurotransmitters.

We next investigated the neural circuit of AgRP neurons. Several AgRP neural circuit have been reported to transmit nutrient signals, such as the AgRP- PVT circuit, which mediates hunger-related food-odor attraction, and the AgRP-BNST circuit, which mediates hunger-induced HPA-axis activation ^29,31^. However, an AgRP circuit specifically involved in amino acid sensing has not been reported. Our work analyzed most AgRP neural circuits (including AgRP to PVT, BNST, DMH, LH, CeA and DR) and surprisingly found that only the circuit of AgRP neurons that project to PVT neurons mediated amino acid appetite. Nevertheless, the AgRP-PVT circuit in glucose or fat sensing needs further investigation and we cannot exclude the roles of other circuits in amino acid sensing under certain conditions. PVT neurons primarily regulate sleep and wakefulness and respond to multiple stresses ^27^. Recent studies have shown that they also regulate feeding and sugar preferences, such as the POMC-PVT circuit, which mediates sugar appetite ^47-49^. However, whether PVT responds to amino acids has not been reported. We found that the AgRP-PVT neural circuit specially regulates amino acid appetite. As PVT neurons can receive projections from AgRP neurons facilitating food seeking ^50^, we assumed that under leucine deprivation, AgRP-PVT projections potentially participate in food seeking for the full amino acid food. Furthermore, combined results from using the antagonist or agonist of respective receptors, we showed that AgRP secreted GABA, but not AgRP or NPY, is the main neurotransmitter that mediates amino acid appetite and acts on PVT neurons. The GABA signal of AgRP mediates HPA axis activation, food intake, and stress-related behaviors to different brain sites ^31,32,51^. We found another role of GABA in amino acid sensing, implying its special function. In PVT, we also found the effect of GABA-B receptor in regulating amino acid preference, which helps in understanding the complex roles of GABA receptors as shown ^52-55^.

When we were looking for the upstream signals acting on AgRP neurons, we focused on the quick detection of amino acids and investigated the function of tongue GCN2 because food first enters the mouth and touches the tongue. Moreover, when excluding visual and olfactory factors, the mice still rapidly detected food containing all EAAs ^56^. This emphasizes the importance of amino acid sensing by the tongue. Though GCN2 is activated in different cells, its function in the tongue has not been investigated. Our results revealed that glossal uncharged leucine tRNA and GCN2-related signals were activated under leucine deficiency, and GCN2 was the main factor that mediates leucine-deprivation-induced amino acid appetite. In our research, we found that the tongue GCN2 knock down resulted in decreased AgRP neural activity under leucine deprivation. Forward and retrograde tracing experiments showed that AgRP neurons received tongue signals through neural connections. Taste signals can be transferred to the brain via taste receptors in the tongue. The signals then act on the nucleus tractus solitarius (NTS), taste cortex, and finally reach the hypothalamus to control glucose utilization, stomach distension, or body weight ^57^. However, no direct connection between AgRP neurons and the tongue has been reported. Our results link tongue signals to the activity and function of AgRP neurons, further providing relevant evidence, however, the accurate circuits between the nutrient, tongue and AgRP neurons require further exploration. Additionally, we cannot exclude other circuits between the tongue and other neurons.

However, several issues remain unsolved in the current study. 1) What is the neural circuit from the tongue to AgRP neurons? Taste neural circuits may be important in this condition, as taste signals can be transmitted to the NTS, which is connected to AgRP neurons. 2) Do other parts of the body contribute to amino acid preference? We cannot exclude the role of other organs or tissues, such as the gut, muscle, and liver, which are important amino acid-sensing organs ^58,59^. These organs may participate in amino acid appetite over longer timescales. For example, fruit flies respond to amino acid deficiency and develop an amino acid appetite through gut GCN2/ATF4 signaling within 2 h ^1^.

In conclusion, this study demonstrated that short-duration EAA deficiency induces rapid and continuous amino acid appetite, whereby quick amino acid detection is mediated by a tongue-AgRP-PVT neural circuit. Tongue GCN2 signaling is necessary for appetite and sends nerve signals to activate AgRP neurons. Together, this work interprets a unique peripheral-to-central neural circuit for sensing amino acid deficiencies, driving amino acid appetite. It expands our understanding of AgRP neuron functions and sheds light on how animals detect and respond to specific nutrients and internal needs.

## Author contributions

S.W., F.Y. and F.G. designed research; F.G., F.Y. and F.X. acquired the funding; S.W., F.Y., Y.G., M.T., S.C., K.T. and P.L. performed animal research; K.L., X.J., H.J., F.X. and W.L. contributed new reagents/analytic tools; S.W., F.Y. and F.G. analyzed data; F.G., F.Y. and S.W. wrote the paper.

## Data, code, and materials availability

All data are available in the main text or the supplementary materials.

## Methods

### Animals

The animal experiments were conducted in accordance with the guidelines of the Institutional Animal Care and Use Committee of Fudan University (Shanghai, China). All mice had a C57BL/6J back ground. AgRP-Cre (*AgRP-irs-Cre*) mice and Ai9 (tdTomato) reporter mice were obtained from Jackson Laboratory (Bar Harbor, ME, USA). POMC-Cre mice were provided by Joel K. Elmquist and Tiemin Liu from the Southwestern Medical Center ^60^. To visualize AgRP or POMC protein-expressing neurons using a fluorescence microscope, AgRP-Cre or POMC-Cre mice were intercrossed with Ai9 (tdTomato) reporter mice. Prior to experiments, mice were group-housed under controlled temperature (23–25 ℃), humidity (50%–60%), and 12-h light/dark cycle (lights on at 8:00 h/lights off at 20:00 h). All mice were provided with water and normal chow diet *ad libitum*, unless otherwise specified. Before commencing the experiments, mice were singly housed for at least 3 days.

### Ethics statement

The conduct of animal research in this study complied with all relevant ethical regulations. The study was approved by the Institutional Animal Care and Use Committee of Fudan University (202203006S).

### Diets and food-choice paradigm

Experimental control (nutritionally complete amino acids), (-) Leu (leucine-deficient), (-) Val (valine-deficient), Ile (isoleucine-deficient), (-) Trp (tryptophan-deficient), (-) Gly (glycine-deficient), and (-) Glu (glutamate-deficient) diets were obtained from Trophic Animal Feed High-Tech Co., Ltd (Nantong, China). The dietary formulations have been previously described ^21^. All diets were isocaloric and compositionally identical in terms of lipid content, with the calorie reduction in the absence of AA compensated by carbohydrates. At the start of the feeding experiment, the mice were acclimated to the control diet for 3 days and then fed the indicated diets. After being fed the control or AA-deficient diets for 3 days (or the indicated period), mice were offered the choice between control versus (-) Leu (or the indicated AA-deficient) diets. For the first 10 min, the bite and touch numbers of the two diets in the mice were counted, and the percentage of the total touch number was calculated. Food intake was measured after 10 and 30 min, and 1, 2, 4, 12, 24 h, or for the indicated time for both diets. The preference index (PI) was calculated as shown below ^1^ (where *f*_c_ is the weight of the control diet consumed by the mice and *f*_d_ is the weight of the AA-deficient diet consumed by the mice).

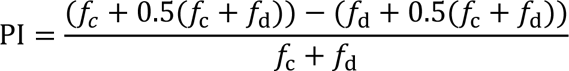

For the fasting experiments, wild-type (WT) mice were acclimated to the control diet for 3 days followed by 0 or 24-h fasting. Then the mice were offered the choice between control versus (-) Leu diets.

For the pair-feeding experiments, WT mice were acclimated to the control diet for 3 days followed by control, (-) Leu, or pair-fed control diet for 3 days, as previously described ^21^. Then the mice were offered the choice between control versus (-) Leu diets.

For the two-bottle solution preference assays, WT mice were acclimated to the control diet for 3 days and then fed the control or (-) Leu diet for 3 days, followed by 24-h thirst. Then the food was withdrawn, and the mice were given a bottle of water and a bottle of 0.75% leucine water solution. The number of licks per bottle was counted during the first 10 min. Water and leucine-containing water intake were measured for 10 min, 30 min, and 1, 2, 3, 4, and 24 h. The PI was calculated as shown below (where *f*_w_ is the weight of water consumed by the mice and *f*_l_ is the weight of leucine solution consumed by the mice consumed).

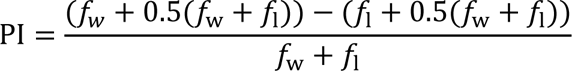

### Stereotaxic surgeries

The mice were anesthetized and placed in a stereotaxic frame (RWD, China). The body temperature of the mice was maintained using a heating pad, and an ophthalmic ointment was applied to maintain eye lubrication. Then, viruses were injected using a microsyringe pump connected to glass pipettes at a rate of 50 nL/min, under the following viral injection coordinates (in mm; midline, bregma, and dorsal surface): for ARC (±0.30 ML, −1.55 AP, −5.80 DV), for aPVT (0 ML, −1.45 AP, −3.05 DV) and pPVT (0 ML, −1.65 AP, −3.05 DV). After injection, the glass pipettes were left in place for 8 min before withdrawal to allow diffusion. The mice were allowed to recover from anesthesia on a heat blanket and then intraperitoneally injected with antibiotics (ceftriaxone sodium, 0.1 g/kg) for 3 days to prevent infection. Mice were group-housed and allowed to recover, while allowing time for the virus to be expressed for at least 3–4 weeks postoperatively.

To knock down VGAT (SLC32A1) in AgRP neurons, AgRP-Cre mice were bilaterally injected with 300 nL Cre-dependent AAV vector, which contained the shSLC32A1 coding sequence and mCherry protein in the opposite orientation, flanked by two inverted loxP sites (AAV2/9-CMV-DIO-shSLC32A1-mCherry; 1.9 × 10^12^ Pfu/mL, HANBIO, Shanghai, China), into the ARC. The target sequence 5’-GATACATTGCACAAGATGAGT-3’ for SLC32A1 was previously validated ^61^. Alternatively, mice were injected with a Cre-dependent AAV vector, containing the scramble vector and mCherry (AAV2/9-CMV-DIO -mCherry; dilute to 1.9× 10^12^ Pfu/mL), as a control. To knock down AGRP and NPY in AgRP neurons, similar strategies were performed. Cre-dependent AAV vectors for AGRP (AAV2/9-Syn-DIO-shAGRP-mCherry; 1 × 10^13^ Pfu/mL, OBiO Tech), NPY (AAV2/9-Syn-DIO-shNPY-mCherry; 1 × 10^13^ Pfu/mL, OBiO Tech, Shanghai, China) or scramble (AAV2/9-Syn-DIO-mCherry; 1 × 10^13^ Pfu/mL, OBiO Tech, Shanghai, China) were injected into the ARC of AgRP-Cre mice. The target sequence for AGRP is 5’-CTGAGTTGTGTTCTGCTGTTG-3’, and the target sequence for NPY is 5’-CGACACTACATCAATCTCA-3’.

### Fiber photometry recording

To measure the calcium signals in AgRP neurons, 300 nL AAV2/9-hSyn-DIO-jGCaMP7s-WPRE-pA virus (1× 10^13^ Pfu/mL, Shanghai Taitool Bioscience Co., Shanghai, China) was injected into the ARC unilaterally in AgRP-Cre mice. To measure the calcium signals in PVT neurons, AAV2/9-mCAMKIIα-jGCaMP7f-WPRE-pA virus (5 × 10^12^ Pfu/mL, Shanghai Taitool Bioscience Co.) was injected into the aPVT and pPVT with a volume of 200 nL per site in WT mice. To measure the GABA signals in PVT neurons, the GABA probe (AAV2/9-hSyn-iGABASnFR-WPRE, 1 × 10^13^ Pfu/mL, Shanghai Taitool Bioscience Co.) was injected into the aPVT and pPVT with a volume of 200 nL per site in WT mice. The optic fiber with 200 μm outer diameter was implanted 100 μm above the viral injection site and fixed to the skull using dental cement powder (C&B, Japan). Following the experiments and histological examination of the brain tissue, the locations of the fiber tips were identified. Four weeks after AAV injection and optic fiber implantation, the mice were subjected to fiber photometry.

The mice were fed a control diet or (-) Leu diet for 3 days, acclimated to the behavioral chamber for more than 20 min prior to recording, and allowed to move freely. Before the experiment, the mice were attached to a patch cable and acclimated for 5 min to stabilize the GCaMP or GABA signals. Then, the signals were recorded for a 5-min baseline and the mice were fed a control diet for 10 min. Photometric data were subjected to minimal processing, consisting of only autofluorescence background subtraction. Blue LED (470 nm) representing the calcium-dependent signal or GABA signal and isosbestic control UV LED (405 nm) fluorescent signals were collected through the recording fiber that was connected to the implant and back to a filtered minicube and into sCMOS cameras, whereby the acquired digital signals were sampled at 1 kHz, demodulated, lock-in amplified, transmitted through a processor, and collected by the provided acquisition software tool.

The data were analyzed using a MATLAB script with minimal processing, consisting of only autofluorescence background subtraction. The transient change in Ca^2+^ was calculated using the function:

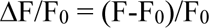

Where F was the raw photometry 470-nm signal and F_0_ was the fitted 405-nm signal. Normalized ΔF/F was calculated by subtracting the baseline ΔF/F from the mean ΔF/F between 0 and 600 s after the commencement of the indicated diet.

### Designer receptors exclusively activated by designer drugs (DREADDs)

To inhibit AgRP neurons with DREADDs, AgRP-Cre mice received stereotaxic injections, bilaterally into the ARC, with a Cre-dependent AAV encoding an inhibitory DREADD GPCR (hM4Di) (AAV2/9-hSyn-DIO-hM4Di-mCherry, 1 × 10^13^ Pfu/mL, OBiO Tech, Shanghai, China), or an AAV encoding only mCherry (AAV2/9-hSyn-DIO-mCherry, 1 × 10^13^ Pfu/mL) as control, at a volume of 300 nL. Four weeks after AAV delivery, all mice received intraperitoneal (i.p.) injections of CNO (MedChemExpress, NJ, USA) at 3 mg/kg body weight for AgRP hM4Di silencing 30 min before the choice assays.

To inhibit neurons with DREADDs, POMC-Cre mice received stereotaxic injections of a Cre-dependent AAV encoding hM4Di (AAV2/9-hSyn-DIO-hM4Di-mCherry, 1 × 10^13^ Pfu/mL, OBiO Tech) or an AAV encoding mCherry (AAV2/9-hSyn-DIO-mCherry, 1 × 10^13^ Pfu/mL) as a control at a volume of 300 nL bilaterally into the ARC. Four weeks after AAV delivery, all mice received i.p. injections of CNO (1.5 mg/kg body weight) for POMC hM4Di silencing 30 min before the choice assay.

To activate PVT glutamatergic neurons with DREADDs, WT mice were administered stereotaxic injections of 200 nL AAVs including a CAMKIIα promoter that encoded hM3Dq (AAV2/9-CAMKIIα-hM3Dq-mCherry, 1.62 × 10^12^ Pfu/mL, OBiO Tech, China), or an AAV encoding only mCherry (AAV2/9-CAMKIIα-mCherry, 1.62 × 10^12^ Pfu/mL, OBiO Tech) as control, respectively into the aPVT and pPVT. To simultaneously inhibit AgRP and PVT glutamatergic neurons using DREADDs, AgRP-Cre mice were administered stereotaxic injections of AAV2/9-hSyn-DIO-hM4Di-mCherry or AAV2/9-hSyn-DIO-mCherry bilaterally into the ARC, and were injected with AAVs including a CAMKIIα promoter encoding hM4Di (AAV2/9-CAMKIIα-hM4Di-mCherry, 5×10^12^ Pfu/mL, OBiO Tech) or AAV2/9-CAMKIIα-mCherry into PVT. Four weeks after AAV delivery, all mice received i.p. injections of CNO at 2 mg/kg for PVT hM3Dq activation or at 3 mg/kg body weight for AgRP/PVT hM4Di silencing.

To inhibit the AgRP innervation of the PVT, AgRP-Cre mice were injected with 200 nL Cre-dependent retroAAV encoding hM4Di (AAV2/2Retro hSyn-DIO-hM4Di-mCherry, 2.21 × 10^13^ Pfu/ml, Shanghai Taitool Bioscience Co.) into the PVT (aPVT and pPVT). Four weeks after AAV delivery, the mice received i.p. injections of saline or 2 mg/kg CNO.

### Optogenetics

To inhibit AgRP neurons with optogenetics, AgRP-Cre mice were injected with a Cre-dependent AAV encoding eNpHR (AAV2/9-EF1-DIO-eNpHR3.0-mCherry, 1 × 10^13^Pfu/mL, OBiO Tech) bilaterally into the ARC. An optical fiber (200 μm) was implant into the third ventricle (0 ML, −1.55 AP, −5.70 DV) as previously reported ^62^. For the circuit of AgRP→PVT, AgRP→DMH, AgRP→BNST, AgRP→CeA, AgRP→DR, AgRP→LH, the mice were injected using AAV2/9-EF1-DIO-eNpHR3.0-mCherry bilaterally into the ARC, with optical fibers (200 μm) respectively implanted into the PVT (0 ML, −1.55 AP, −2.95 DV), DMH (0 ML, −1.55 AP, −5.0 DV), BNST (±1.4 ML, +0.62 AP, −4.15 DV, with 10° ML angle), CeA (±3.0 ML, −1.42 AP, −4.7 DV), DR (−1.05 ML, −4.3 AP, −2.7 DV, with 20° ML angle) and LH (±2.1 ML, -0.94 AP, –4.71 DV, with 10° ML angle). Before the choice assays, the mice were fed control diets for 3 days and then (-) Leu diets for 3 days, followed by 24-h fasting to allow the mice to eat food in a short time.

The mice were subjected to persistent photoinhibition (yellow light, 589 nm, 15 mW for AgRP neurons, 8 mW for AgRP-other site circuits, constant) for 5 min without food. Then the mice were given control and (-) Leu food with the light on for 10 min. The number of touches on each food pellet and preference index within 10 min were recorded and calculated.

### Brain cannulation and drug injections

A cannula was placed into the PVT (coordinates: ML: 0 mm, AP: −1.5 mm, DV: −3.0 mm from bregma) or third ventricle (ML: 0 mm, AP: −1.55 mm, DV: −5.5 mm from bregma). Two screws were placed in the lambdoid structure to aid in supporting the cannula in the skull with dental cement. The mice were allowed to recover from anesthesia on a heat blanket and were injected for 3 days with antibiotics (ceftriaxone sodium, 0.1 g/kg i.p.) to prevent infection. The mice were housed individually and allowed to recover for 10 days after surgery. Start from the (-) Leu diet feeding, the drugs (1 μL) were injected via cannula daily. On the day of the choice assays, the mice received drugs via a cannula 30 min before food was provided. The drugs including GABAbR antagonist saclofen (0.1 μg/μL, HY-100813 MCE) and CGP55845 hydrochloride (8 ng/μL, HY-103516, MCE, China), GABAbR agonist baclofen (0.04 μg/μL, HY-B0007 MCE), GABAaR antagonist bicuculline (2.5 μM, HY-N0219, MCE), MC3R/4R antagonist SHU 9119 (1 nM, HY-P0227, MCE), NPY1R antagonist BIBO3304 (0.25 mM, HY-107725, MCE), NPY2R antagonist BIIE-0246 (0.25 mM, HY-101986), NPY5R antagonist CGP71683 hydrochloride(15 nM, HY-107723, MCE) and leucine (1.1μg/μL in 10% DMSO).

### Tongue AAV injection

To knock down tongue GCN2 in WT or AgRP/Ai9 mice, the mice received injections into the tongue of AAV2/DJ-shGCN2-EGFP (2 × 10^11^ Pfu/mL, OBiO Tech) or AAV2/DJ-EGFP (2 × 10^11^ Pfu/mL, OBiO Tech), as a control group, as previously described ^63^. The target sequence 5ʹ-TCTGGATGGATTAGCTTATA-3ʹ for GCN2 was previously validated ^64^. Specifically, the mouse tongue was pierced parallel to the tongue epithelium with an insulin needle from the tip to the middle and posterior parts of the tongue to create a channel between the epithelium and muscle layer. A 25-μL microsyringe was injected along the channel into the middle and posterior part of the tongue. The syringe was slowly withdrawn while injecting the virus, and the direction of the needle tip was changed to ensure that the virus evenly infected different parts of the tongue. Inject 2–3 μL virus at each site, to a total volume of 25 μL. Finally, the wound was sealed with tissue adhesive to prevent viral outflow. Three weeks later, we dropped virus to the surface of the tongue, at a rate of 2 μL each time, and repeated this after the virus was absorbed, up to a total volume of 30 μL.

To knock down tongue GCN2 and activate AgRP neurons, AgRP-Cre mice received injections into the tongue of AAV2/DJ-shGCN2-EGFP (2×10^11^ Pfu/mL, OBiO Tech), and injections bilaterally into the ARC of AAV2/9-hSyn-DIO-hM3Dq-mCherry (5×10^12^ Pfu/mL, OBiO Tech) or AAV2/9-hSyn-DIO-mCherry (5×10^12^ Pfu/mL, OBiO Tech), using above-described methods.

### PRV and HSV injections

WT male mice were anesthetized and injected with pseudorabies virus (BC-PRV-mCherry, 5 × 10^9^ Pfu/mL, BrainCase Co., Ltd, China) into the ARC, and herpes simplex virus (BC-HSV-EGFP, 1 × 10^8^ Pfu/mL, BrainCase Co., Ltd, China) into the tongue.

AgRP-Cre male mice were anesthetized and injected with pseudorabies virus (BC-PRV-DIO-CAG-mCherry, 5 × 10^9^ Pfu/mL, BrainCase Co., Ltd, China) into the ARC, and herpes simplex virus (BC-HSV-EGFP, 1 × 10^8^ Pfu/mL, BrainCase Co., Ltd, China) into the tongue. After 3–5 days, the brains and tongues were collected.

### IF staining

The mice were transcardially perfused with saline followed by phosphate-buffered saline (PBS) containing 4% paraformaldehyde (PFA). Mouse brains were dissected and fixed overnight at 4 °C in 4% PFA, followed by cryoprotection in PBS containing 20% and 30% sucrose at 4 °C. Free-floating sections (25 μm) were prepared with a cryostat. Slices were blocked for 1 h at room temperature in PBST (0.3% Triton X-100) with 5% normal donkey serum, followed by incubation with primary antibodies at 4 °C overnight and secondary antibodies at room temperature for 2 h. Primary antibodies used in IF experiments included anti-VGAT (1:2,000, ZRB1158, Sigma), anti-mCherry (1:500, ARG55723, Arigobio) and anti-c-Fos (1:1,000, #2250, Cell Signaling Technology).

### Measurement of serum amino acids

Mouse serum was collected, added to four volumes of organic reagent (methanol: acetonitrile = 1:1), and vortexed for 30 s. Then, the liquid was treated with ultrasonication to thoroughly disrupt the cells and then allowed to subside for 1 h at 20 °C. The liquid was centrifuged at 12,000 rpm for 15 min, and the supernatant was evaporated to dryness. For the residue, 100 μL 10% acetonitrile was added and then mixed for measurement. Serum amino acids were analyzed using high-performance liquid chromatography (Ultimate 3000, USA) and tandem mass spectrometry (API 3200 Q-TRAP, USA) by Beijing MS Medical Research Co. Ltd. (Beijing, China).

### Western blot

Tissues were homogenized in ice-cold lysis buffer (50 mM Tris HCl, pH 7.5, 0.5% Nonidet P-40, 150 mM NaCl, 2 mM EGTA, 1 mM Na_3_VO_4_, 100 mM NaF, 10 mM Na_4_P_2_O_7_, 1 mM phenylmethylsulfonyl fluoride, 10 μg/mL aprotinin, 10 μg/mL leupeptin). Tissue extracts were then immunoblotted with anti-p-eIf2α (1:1,000, #3398, Cell Signaling Technology), anti-t-eIf2α (1:1,000, #9722, Cell Signaling Technology) and anti-α-tubulin (1:2,000, 66031-1-AP, Proteintech, China) primary antibodies.

### Uncharged tRNA

Total and uncharged tRNA levels were measured using the four-leaf clover quantitative real-time PCR (qRT-PCR) method as previously described ^65,66^. Briefly, RNA was extracted from the tongue tissue of mice under control or leucine-deficient diet treatment using TRIzol and dissolved in 0.1 M NaAC [pH 5.2]. RNA was treated with either NaIO_4_, which oxidizes the 3’-A residue of uncharged tRNAs, but not of charged tRNA, owing to the protection of amino acids, or NaCl as a parallel mock treatment in the dark at room temperature for 20 min, followed by quenching with 0.3 M glucose for 15 min. Then, the samples were desalinated and discharged (deacylated) in a solution of 100 mM Tris-HCl [pH 9.0] at 37 °C for 45 min. Following deacylation, DNA/RNA-hybrid adaptor (5’-/Phos/TCGTAGGGTCCGAG GTATTCACGATGrGrCrN-3’) was ligated to tRNAs using T4 RNA ligase 2 (NEB). Next, an oligo (5’-ATACCTCGGACCCTACGA-3’), complementary to the adaptor sequence, was used for cDNA synthesis using HiScript III RT kit (Vazyme). cDNA was used for the qPCR-based detection of tRNAs. The primers used for qRT-PCR are listed in **Supplementary Table 1** (*LeuWAG*).

### RNA isolation and RT-PCR

RNA was extracted using the TRIzol reagent (Invitrogen). The mRNA was reverse-transcribed using a High-Capacity cDNA Reverse Transcription Kit (Thermo Scientific) and subjected to quantitative real-time PCR analysis using SYBR Green I Master Mix reagent on an ABI 7900 system (Applied Biosystems). Primer sequences used in this study are listed in **Supplementary Table 1**.

### Statistical analysis

Statistical analyses were performed using GraphPad Prism software (version 8.0; GraphPad Software, San Diego, CA, USA). All values are presented as the mean ± standard error of the mean (SEM). A two-tailed unpaired Student’s *t*-test was used to compare the two groups. For experiments involving multiple comparisons, data were analyzed using one-way analysis of variance (ANOVA) followed by Dunnett’s/Tukey’s multiple comparison test, or two-way ANOVA followed by Tukey’s multiple comparison test. Individual data points on every histogram reflect the individual variability of the measures. Statistical significance was defined as *\*P* < 0.05, *\*\*P* < 0.01, *\*\*\*P* < 0.001, *\*\*\*\*P* < 0.0001. Sample sizes and *P*-values are shown in the figure legends.

