## Extended Data Fig. 1-9; Supplementary Table 1 for "The tongue-brain axis mediates a hidden amino acid appetite"

Address: 131 Dong' an Road Road, Shanghai, China 200032

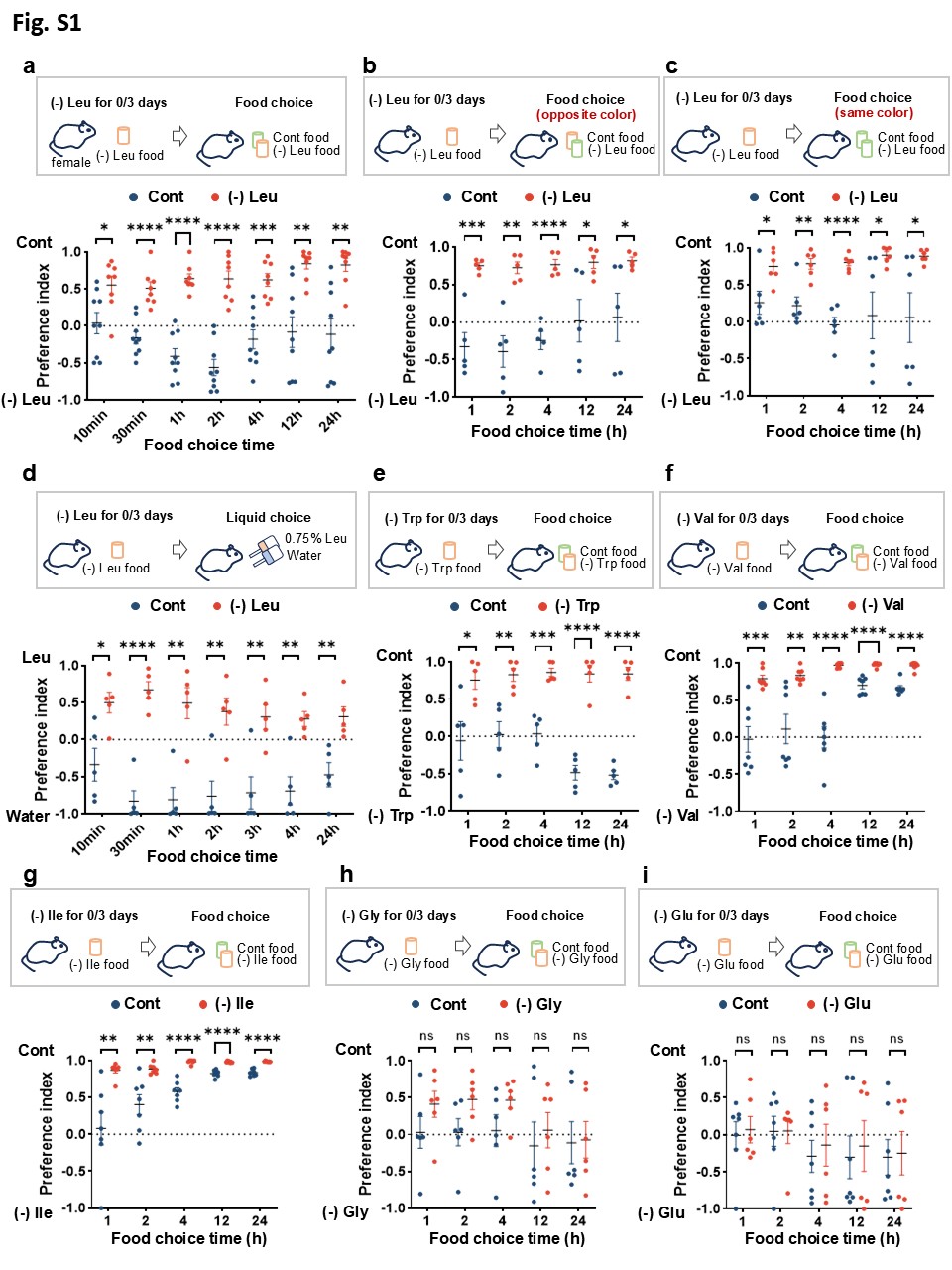

**Extended Data Fig. 1 Essential amino acid deprivation induce amino acid appetite under different situations**

a, Two-choice preferences for indicated time (10min, 30min, 1h, 2h, 4h, 12h and 24h) of female mice that fed a control (Cont) or leucine-deficient [(-) Leu] diet for 3 days.

b, Two-choice preferences using the opposite color of the food for indicated time (1h, 2h, 4h, 12h and 24h) of mice that fed a Cont or (-) Leu diet for 3 days.

c, Two-choice preferences using the same color of the food for indicated time (1h, 2h, 4h, 12h and 24h) of mice that fed a Cont or (-) Leu diet for 3 days.

d, Two-choice preferences between leucine solution and water for indicated time (10min, 30min, 1h, 2h, 4h, 12h and 24h) of mice that fed a Cont or (-) Leu diet for 3 days.

e, Two-choice preferences for indicated time (1h, 2h, 4h, 12h and 24h) of mice that fed a Cont or tryptophan-deficient [(-) Trp] diet for 3 days.

f, Two-choice preferences for indicated time (1h, 2h, 4h, 12h and 24h) of mice that fed a Cont or valine-deficient [(-) Val] diet for 3 days.

g, Two-choice preferences for indicated time (1h, 2h, 4h, 12h and 24h) of mice that fed a Cont or isoleucine-deficient [(-) Ile] diet for 3 days.

h, Two-choice preferences for indicated time (1h, 2h, 4h, 12h and 24h) of mice that fed a Cont or glycine-deficient [(-) Gly] diet for 3 days.

i, Two-choice preferences for indicated time (1h, 2h, 4h, 12h and 24h) of mice that fed a control or glutamate-deficient [(-) Glu] diet for 3 days.

Studies for a were conducted using 8–10-week-old female wild-type (WT) mice fed a Cont or (-) Leu diet for 3 days; studies for b to i were conducted using 8–10-week-old male WT mice fed a Cont, (-) Leu, (-) Trp, (-) Val, (-) Ile, (-) Gly, or (-) Glu diet for 3 days. Data are expressed as the mean ± SEM (n = 5-9 per group, as indicated), with individual data points. Data were analyzed via two-tailed unpaired Student’s t-test. *P < 0.05, **P < 0.01, ***P < 0.001, ****P < 0.0001.

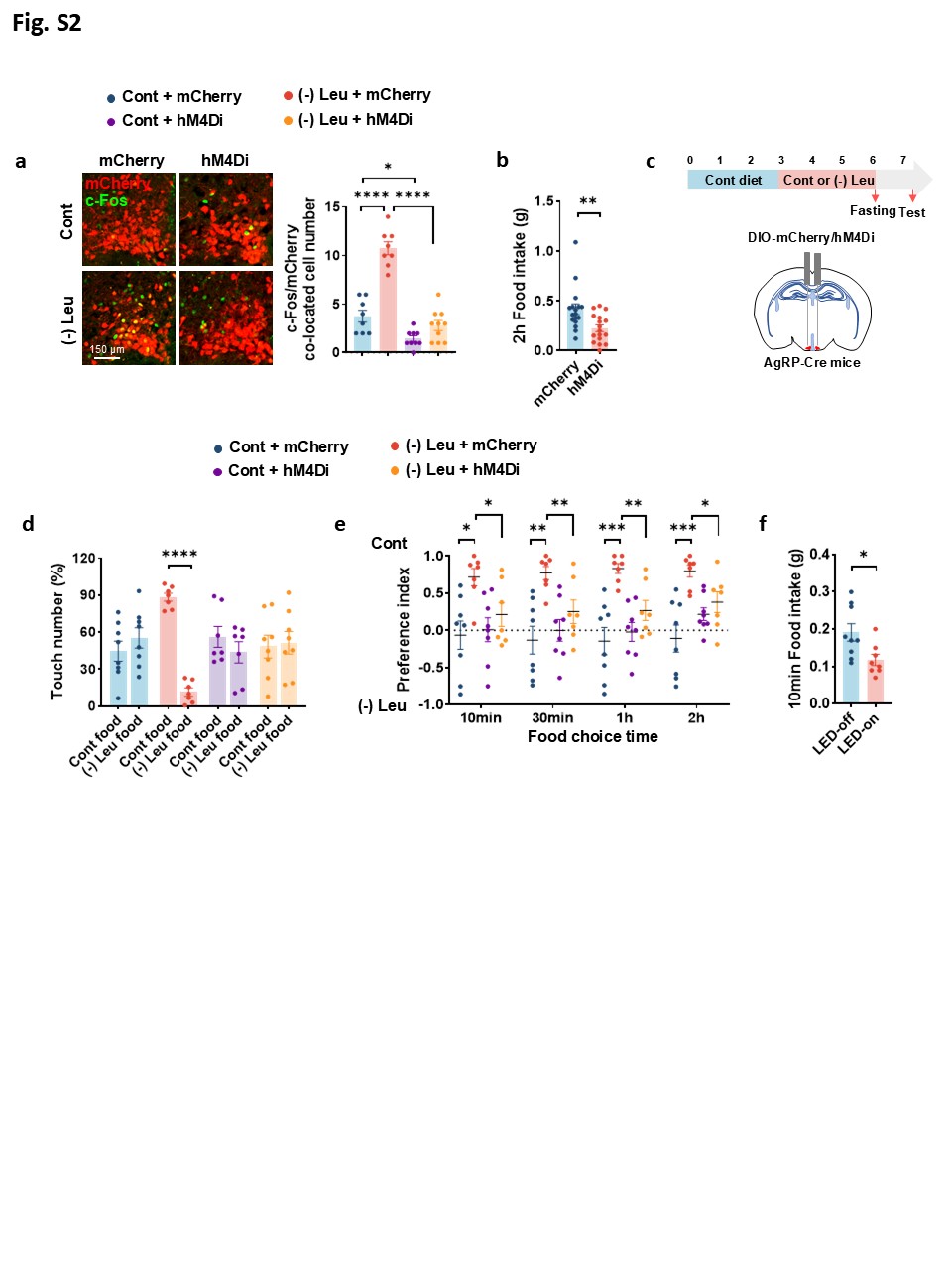

**Extended Data Fig. 2 Inhibition of hypothalamic AgRP neurons reduced feeding and abolished leucine deprivation-induced amino acid appetite.**

a, Immunofluorescence (IF) staining for mCherry (red), c-Fos (green) or merge (yellow) in ARC (left), and quantification of c-Fos and mCherry colocalized cell numbers (right).

b, 2-h food intake after CNO injection.

c, Schematics illustrating virus-mediated mCherry/hM4Di expression (red) in ARC^AgRP^ neurons and the timeline of diet feeding as well as the test. Mice were fed a control (Cont) or leucine-deficient [(-) Leu] diet for 3 days followed by fasting 24h and CNO injection.

d, Touch number percentage for the first 10 min of two-choice assays in c.

e, Two-choice preferences for indicated time (10min, 30min, 1h, 2h) in c.

f, 10-min food intake with or without yellow light.

Studies for a to e were conducted using 12-16-week-old male AgRP-Cre mice receiving AAVs expressing mCherry or hM4Di fed a Cont or (-) Leu diet for 3 days; studies for f were conducted using 12-16-week-old male AgRP-Cre mice receiving AAVs expressing NpHR (AAV2/9-EF1-DIO-eNpHR3.0-mCherry). Data are expressed as the mean ± SEM (n = 7-17 per group, as indicated), with individual data points. Data were analyzed via two-tailed unpaired Student’s t-test (b, d, f), or two-way ANOVA followed by Tukey’s multiple comparisons test (a, e). *P < 0.05, **P < 0.01, ***P < 0.001, ****P < 0.0001.

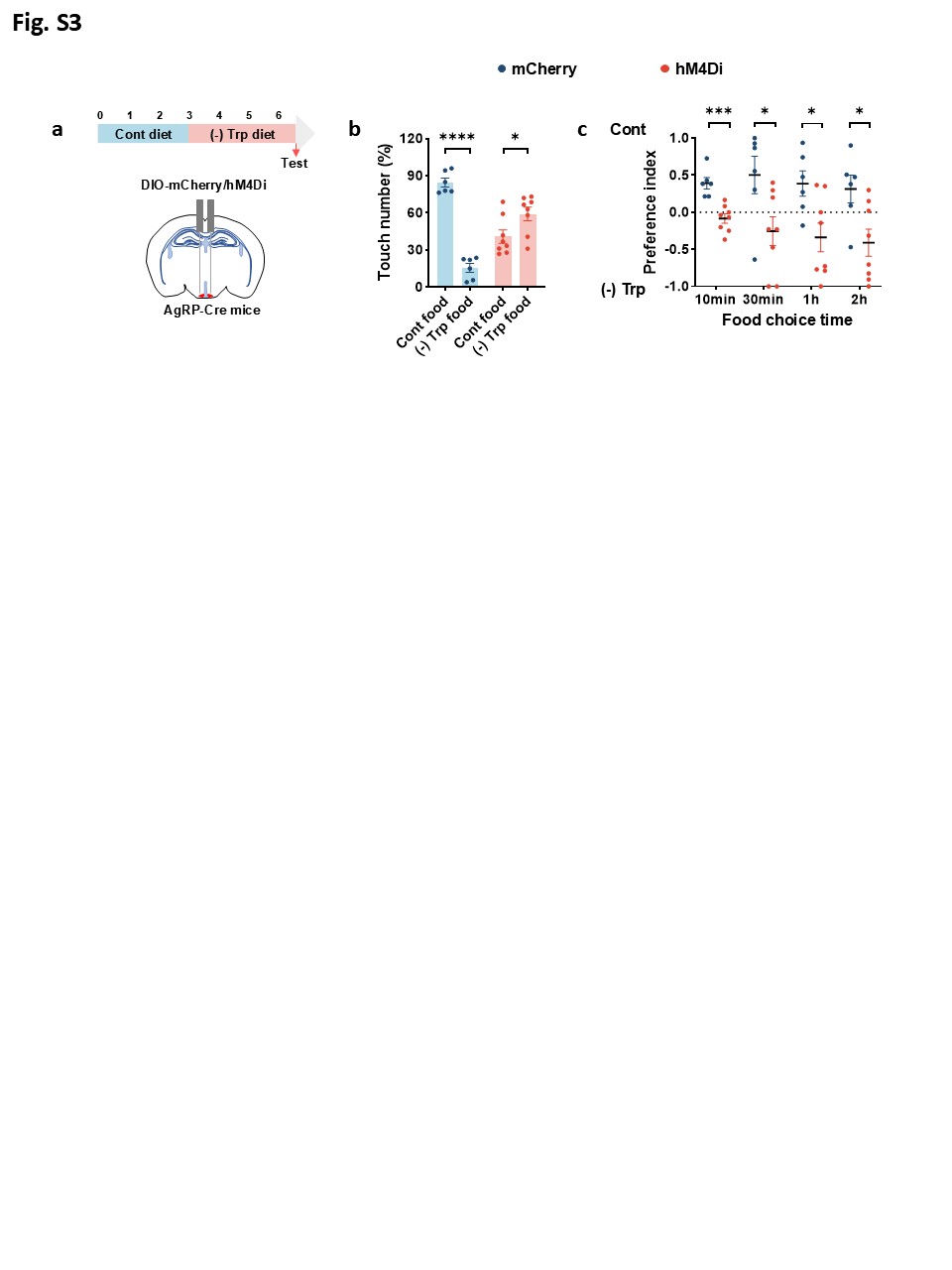

**Extended Data Fig. 3 Inhibition of AgRP neurons abolished tryptophan deprivation-induced amino acid appetite**

a, Schematics illustrating virus-mediated mCherry/hM4Di expression (red) in ARC^AgRP^ neurons and the timeline of diet feeding as well as the test. Mice were fed a tryptophan-deficient [(-) Trp] diet for 3 days followed by CNO injection.

b, Touch number percentage for the first 10 min of two-choice assays in a.

c, Two-choice preferences for indicated time (10min, 30min, 1h, 2h) in a.

Studies were conducted using 12-16-week-old male AgRP-Cre mice receiving AAVs expressing mCherry or hM4Di fed a (-) Trp diet for 3 days. Data are expressed as the mean ± SEM (n = 6-8 per group, as indicated), with individual data points. Data were analyzed via two-tailed unpaired Student’s t-test. *P < 0.05, ***P < 0.001, ****P < 0.0001.

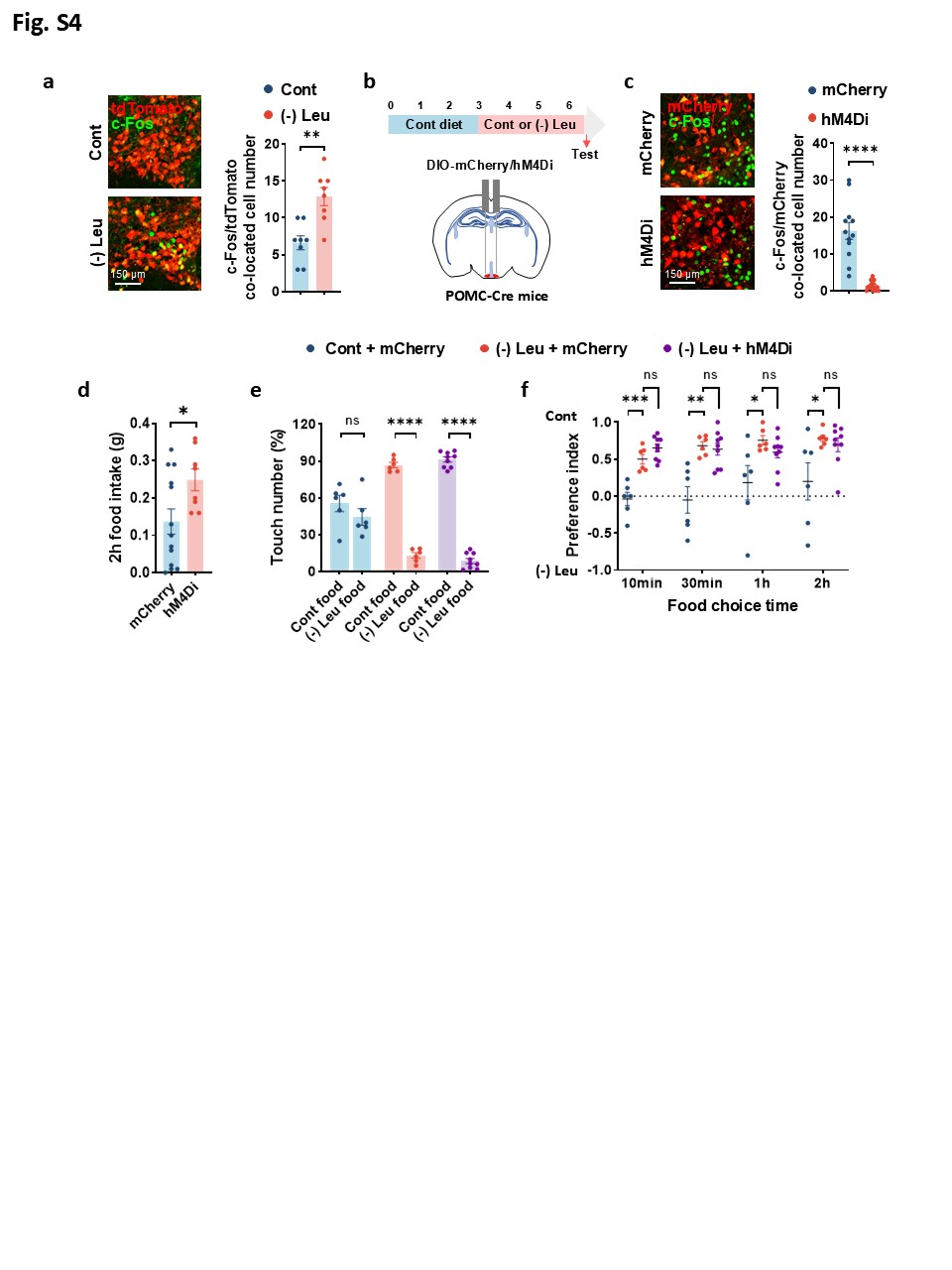

**Extended Data Fig. 4 Inhibition of POMC neurons had no impact on leucine deprivation-induced amino acid appetite**

a, Immunofluorescence (IF) staining for tdTomato (red), c-Fos (green) or merge (yellow) in arcuate nucleus (ARC) (left), and quantification of c-Fos and tdTomato colocalized cell numbers (right).

b, Schematics illustrating virus-mediated mCherry/hM4Di expression (red) in ARC^POMC^ neurons and the timeline of diet feeding as well as the test. Mice were fed a control (Cont) or leucine-deficient [(-) Leu] diet for 3 days followed by CNO injection.

c, IF staining for mCherry (red), c-Fos (green) or merge (yellow) in ARC (left), and quantification of c-Fos and mCherry colocalized cell numbers (right).

d, 2-h food intake after CNO injection.

e, Touch number percentage for the first 10 min of two-choice assays in b.

f, Two-choice preferences for indicated time (10min, 30min, 1h, 2h) in b.

Studies for a were conducted using 40-41-week-old male POMC/Ai9 mice fed a Cont or (-) Leu diet for 3 days; studies for b to f were conducted using 10-12-week-old male POMC-Cre mice receiving AAVs expressing mCherry or hM4Di fed a control or leucine-deficient diet for 3 days. Data are expressed as the mean ± SEM (n = 6-15 per group, as indicated), with individual data points. Data were analyzed via two-tailed unpaired Student’s t-test (a, c, d, e), or one-way ANOVA followed by Tukey’ s multiple comparisons test f. *P < 0.05, **P < 0.01, ***P < 0.001, ****P < 0.0001.

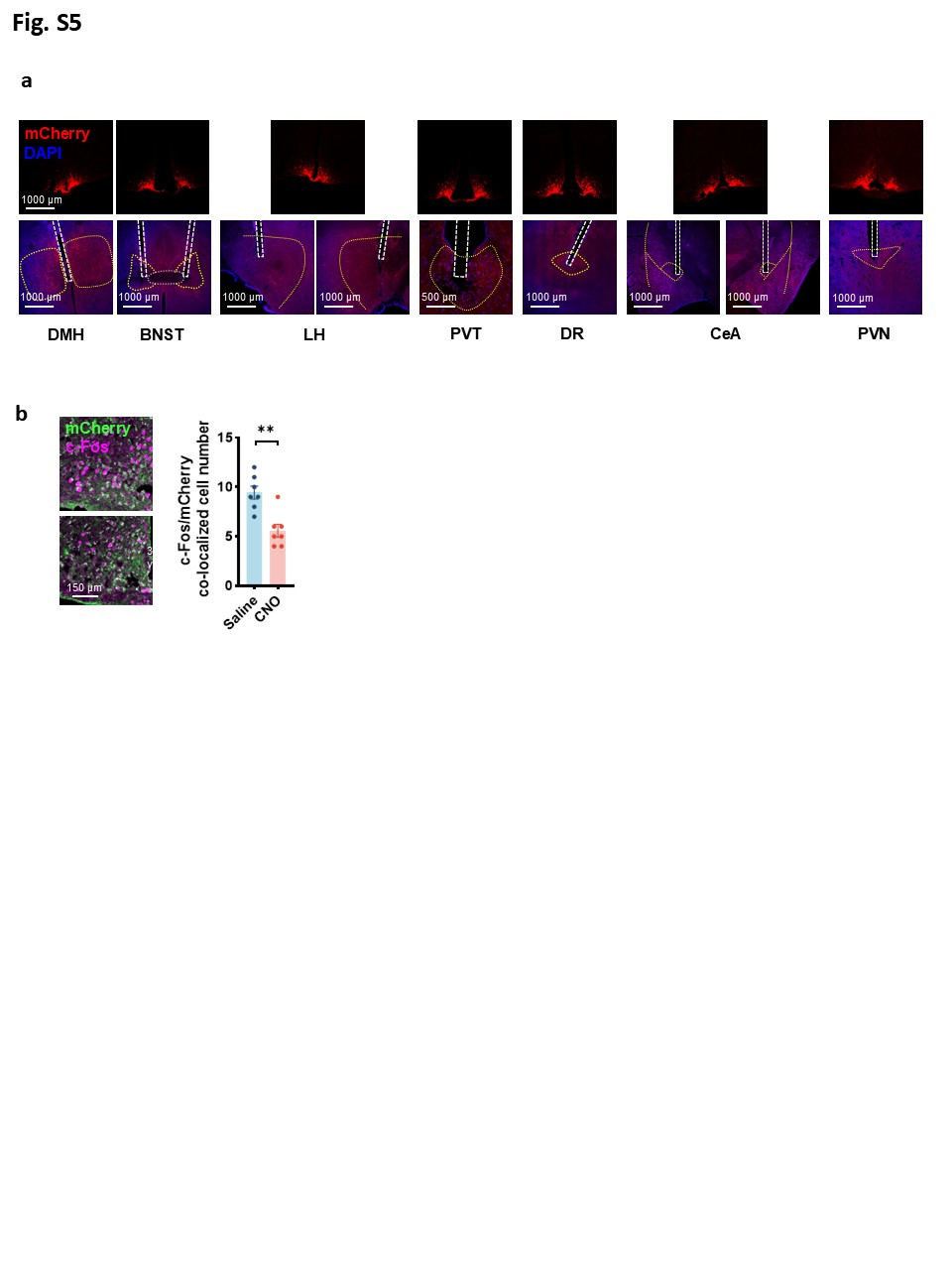

**Extended Data Fig. 5 Validation of the effectiveness of the virus and the fibre locations**

a, Immunofluorescence (IF) staining for mCherry (red), DAPI (blue) in arcuate nucleus (ARC) (top) and the fibre locations (bottom) in dorsomedial hypothalamic nucleus (DMH), bed nucleus of the stria terminalis (BNST), lateral hypothalamus (LH), paraventricular thalamus (PVT), dorsa raphe (DR), central amygdala (CeA) and hypothalamic paraventricular nucleus (PVN).

b, IF staining for for mCherry (green), c-Fos (magenta) or merge (white) in ARC (left), and quantification of c-Fos and mCherry colocalized cell numbers (right); 3V, third ventricle.

Studies for a were conducted using 8-14-week-old male AgRP-Cre mice (DMH, BNST, PVT and DR), or using 12-16-week-old female AgRP-Cre mice (LH, CeA, PVN) receiving AAVs expressing NpHR (AAV2/9-EF1-DIO-eNpHR3.0-mCherry) with an optical fiber implantation into the indicated locations, fed a leucine-deficient [(-) Leu] diet for 3 days; studies for b were conducted using 30-32-week-old male AgRP-Cre mice receiving AAVs expressing Retro-DIO-hM4Di-mCherry fed a (-) Leu diet for 3 days. Data are expressed as the mean ± SEM (n = 7 per group, as indicated), with individual data points. Data were analyzed via two-tailed unpaired Student’s t-test. **P < 0.01.

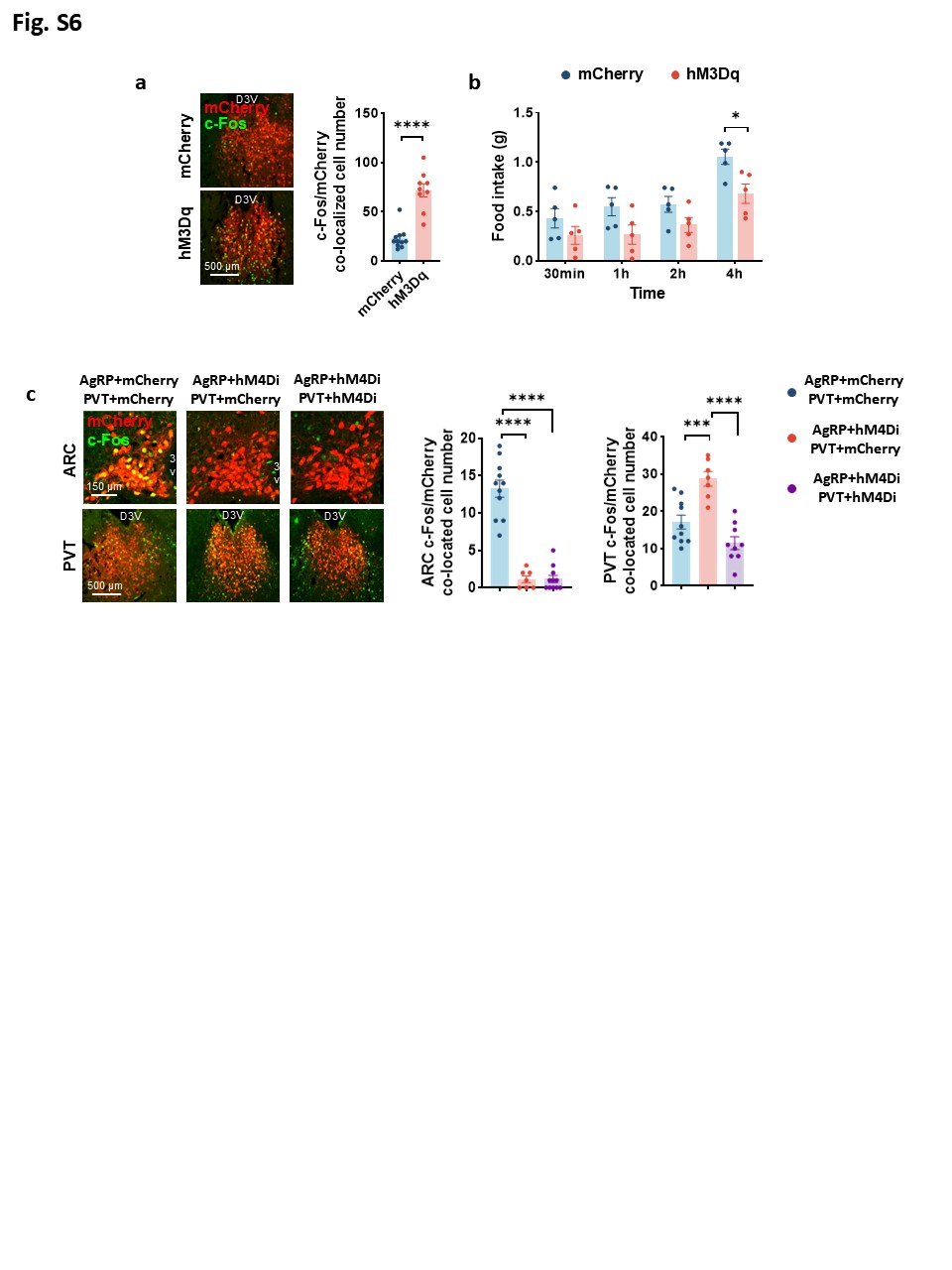

**Extended Data Fig. 6 Validation of the effectiveness of the PVT^Glu^ neural activation and AgRP neural inhibition**

a, Immunofluorescence (IF) staining for mCherry (red), c-Fos (green) or merge (yellow) in paraventricular thalamus (PVT, left), and quantification of c-Fos and mCherry colocalized cell numbers (right); D3V, dorsal third ventricle.

b, Food intake after CNO injection.

c, IF staining for mCherry (red), c-Fos (green) or merge (yellow) in ARC, and for mCherry (red), c-Fos (green) or merge (yellow) in PVT (left), and quantification of c-Fos cell numbers (right); 3V, third ventricle. Studies for a and b were conducted using 10-12-week-old male WT mice receiving AAVs expressing mCherry or hM3Dq; studies for c were conducted using 12-18-week-old male AgRP-Cre mice receiving AAVs expressing DIO-mCherry or DIO-hM4Di in ARC, and CaMKIIa-mCherry or CaMKIIa-hM4Di in PVT, fed a leucine-deficient diet for 3 days. Data are expressed as the mean ± SEM (n = 5-11 per group, as indicated), with individual data points. Data were analyzed via two-tailed unpaired Student’s t-test (a, b), or one-way ANOVA followed by Dunnett’s multiple comparisons test (c, left), or one-way ANOVA followed by Tukey’s multiple comparisons test (c, right). ***P < 0.001, ****P < 0.0001.

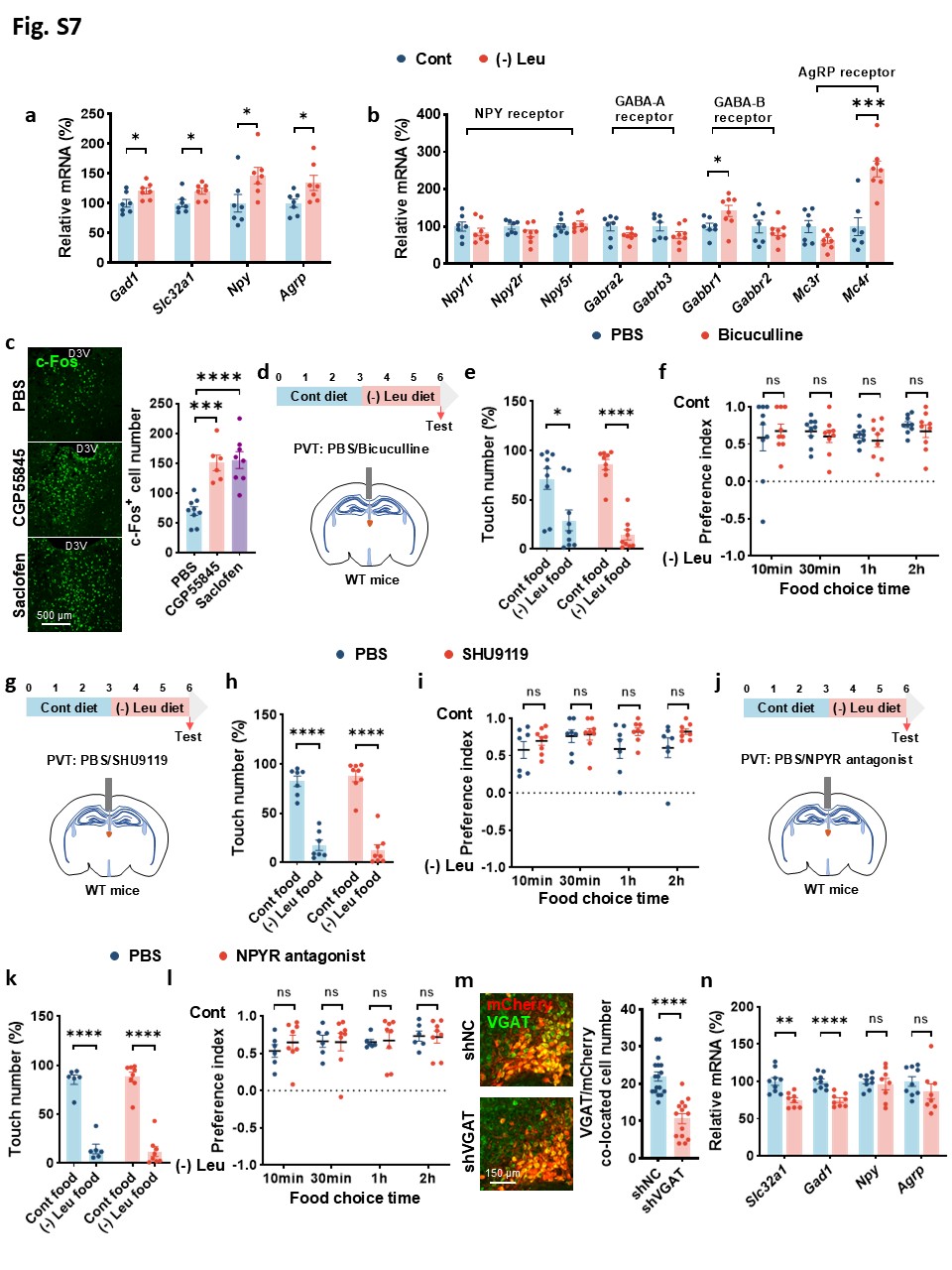

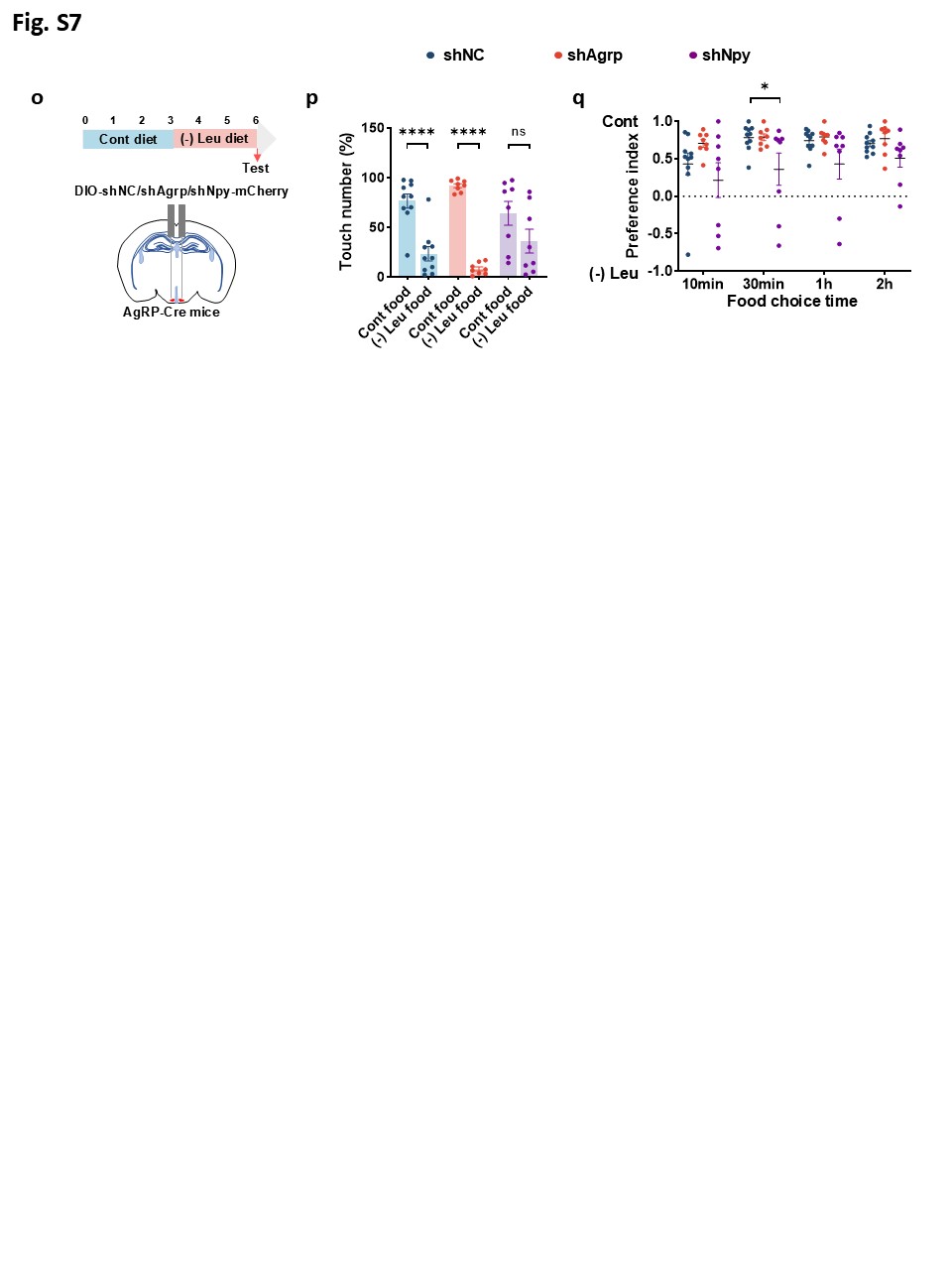

**Extended Data Fig. 7 AgRP neurons act to PVT^Glu^ neurons via GABA signaling, but not other neurotransmitters, mediating amino acid appetite**

a, Gene expression of *Gad1*, *Slc32a1*, *Npy* and *Agrp* in hypothalamic arcuate nucleus (ARC) by RT-PCR.

b, Gene expression of *Npy1r*, *Npy2r*, *Npy5r*, *Gabra2*, *Gabrb3*, *Gabbr1*, *Gabbr2*, *Mc3r* and *Mc4r* in paraventricular thalamus (PVT) by RT-PCR.

c, Immunofluorescence (IF) staining for c-Fos (green) in PVT (left), and quantification of c-Fos cell numbers (right); D3V, dorsal third ventricle.

d, Schematics illustrating PVT injection of Bicuculline or PBS, and the timeline of diet feeding as well as the test. Mice were fed a leucine-deficient [(-) Leu] diet for 3 days followed by PVT injection.

e, Touch number percentage for the first 10 min of two-choice assays in d.

f, Two-choice preferences for indicated time (10min, 30min, 1h, 2h) in d.

g, Schematics illustrating PVT injection of SHU9119 or PBS, and the timeline of diet feeding as well as the test. Mice were fed a (-) Leu diet for 3 days followed by PVT injection.

h, Touch number percentage for the first 10 min of two-choice assays in g.

i, Two-choice preferences for indicated time (10min, 30min, 1h, 2h) in g.

j, Schematics illustrating PVT injection of NPYR antagonists or PBS, and the timeline of diet feeding as well as the test. Mice were fed a (-) Leu diet for 3 days followed by PVT injection.

k, Touch number percentage for the first 10 min of two-choice assays in j.

l, Two-choice preferences for indicated time (10min, 30min, 1h, 2h) in j.

m, IF staining for mCherry (red), VGAT (Green) or merge (yellow) in ARC (left), and quantification of VGAT and mCherry colocalized cell numbers (right).

n, Gene expression of *Gad1*, *Slc32a1*, *Npy* and *Agrp* in hypothalamic arcuate nucleus (ARC) by RT-PCR.

o, Schematics illustrating virus-mediated shNC, shAgRP or shNpy expression (red) in ARC^AgRP^ neurons and the timeline of diet feeding as well as the test. Mice were fed a (-) Leu diet for 3 days.

p, Touch number percentage for the first 10 min of two-choice assays in o.

q, Two-choice preferences for indicated time (10min, 30min, 1h, 2h) in o.

Studies for a and b were conducted using 8–10-week-old male wild-type (WT) mice fed a control or (-) Leu diet for 3 days; studies for c were conducted using 10-12-week-old male WT mice with PVT injection of Saclofen, CGP55845 or PBS, fed a (-) Leu diet for 3 days; studies for d to l were conducted using 10-12-week-old male WT mice with PVT injection of Bicuculline, SHU9119, NPYR antagonists or PBS, fed a (-) Leu diet for 3 days; studies for m and n were conducted using 8-12-week-old female AgRP-Cre mice receiving AAVs expressing DIO-shNC or DIO-shVGAT in ARC fed a (-) Leu diet for 3 days; studies for o-q were conducted using 15-16-week-old male AgRP-Cre mice receiving AAVs expressing DIO-shNC, DIO-shAgRP or DIO-shNpy in ARC fed a (-) Leu diet for 3 days. Data are expressed as the mean ± SEM (n = 6-17 per group, as indicated), with individual data points. Data were analyzed via two-tailed unpaired Student’s t-test (a, b, e, f, h, i, k, l, m, n, p), or one-way ANOVA followed by Dunnett’s multiple comparisons test (c, q). *P < 0.05, **P < 0.01, ***P < 0.001, ****P < 0.0001.

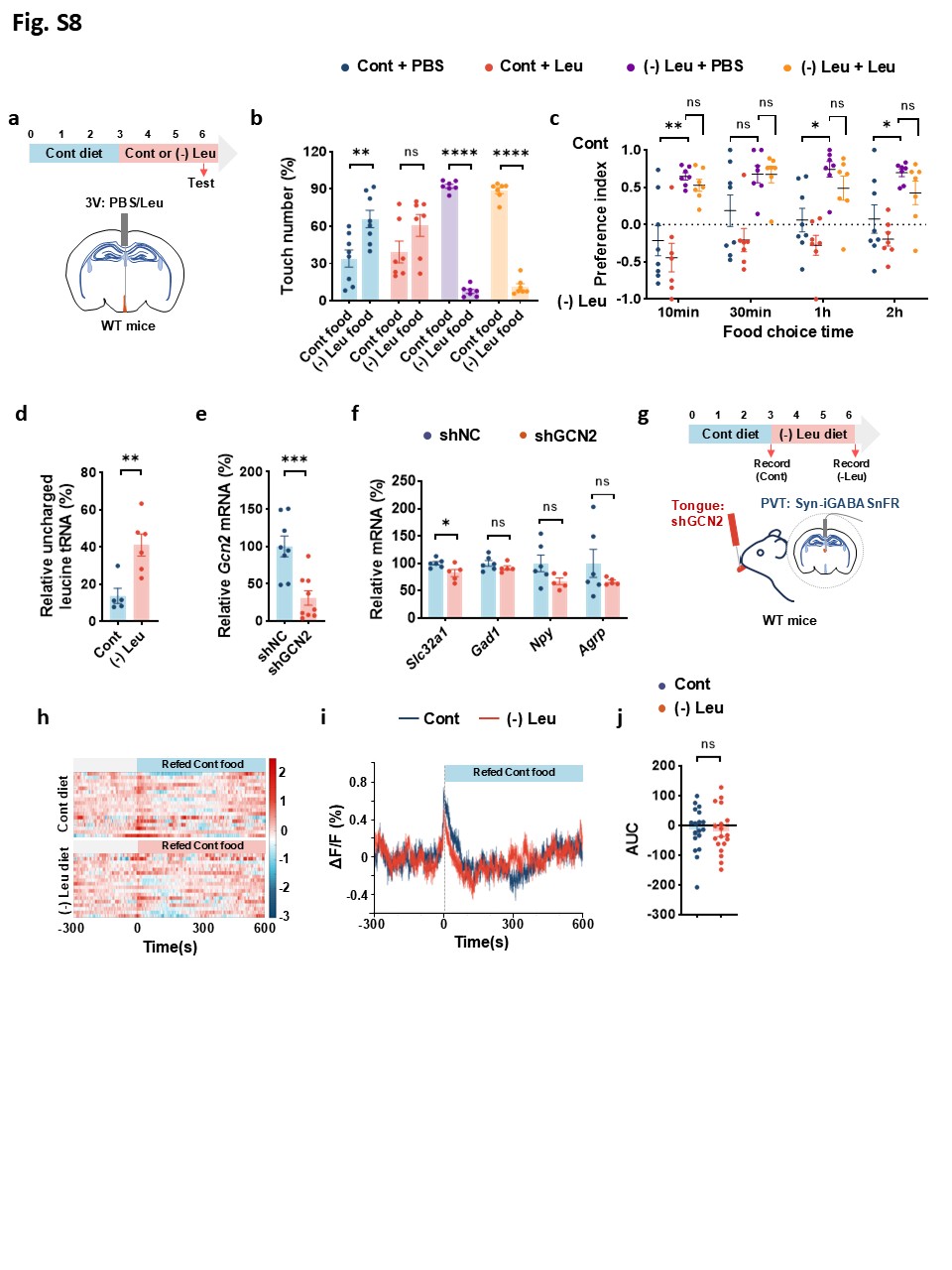

**Extended Data Fig. 8 The tongue GCN2, not brain leucine levels, mediates leucine deprivation-induced amino acid appetite**

a, Schematics illustrating intracerebroventricular (ICV) injection of leucine or PBS, and the timeline of diet feeding as well as the test. Mice were fed a control (Cont) or leucine-deficient [(-) Leu] diet for 3 days followed by ICV injection.

b, Touch number percentage for the first 10 min of two-choice assays in a.

c, Two-choice preferences for indicated time (10min, 30min, 1h, 2h) in a.

d, Relative levels of uncharged tRNA for leucine by RT-PCR.

e, Gene expression of *Gcn2* in the tongue by RT-PCR.

f, Gene expression of *Slc32a1*, *Gad1*, *Npy* and *Agrp* in ARC by RT-PCR.

g, Schematics illustrating virus-mediated shGCN2 expression (red) in the tongue, the fiber photometry recordings of GABA signals in PVT neurons, and the timeline of diet feeding as well as the test. Each mouse was subjected to the experiment twice.

h, Heatmap of GABA signals in PVT neurons of mice that fed a Cont or (-) Leu diet for 3 days in response to a Cont diet. Each heatmap represents a single behavioral session.

i, Averaged traces of GABA signals in h.

j, The area under curve (AUC) of the GABA signals (0-600s) in i.

Studies for a to c were conducted using 10-12-week-old male wild-type (WT) mice with ICV injection of leucine or PBS, fed a Cont or (-) Leu diet for 3 days; studies for d were conducted using 8-12-week-old male WT mice fed a Cont or (-) Leu diet for 3 days; studies for e and f were conducted using 8-10-week-old male WT mice receiving AAVs expressing shGCN2 or shNC in the tongue; studies for g-j were conducted using 10-12-week-old male WT mice receiving AAVs expressing shGCN2 in the tongue, and Syn-iGABASnFR in PVT, fed a Cont or (-) Leu diet for 3 days. Data are expressed as the mean ± SEM (n = 5-18 per group, as indicated), with individual data points. Data were analyzed via two-way ANOVA followed by Tukey’s multiple comparisons test c, or via two-tailed unpaired Student’s t-test (b, d, e, f, j). *P < 0.05, **P < 0.01, ***P < 0.001, ****P < 0.0001.

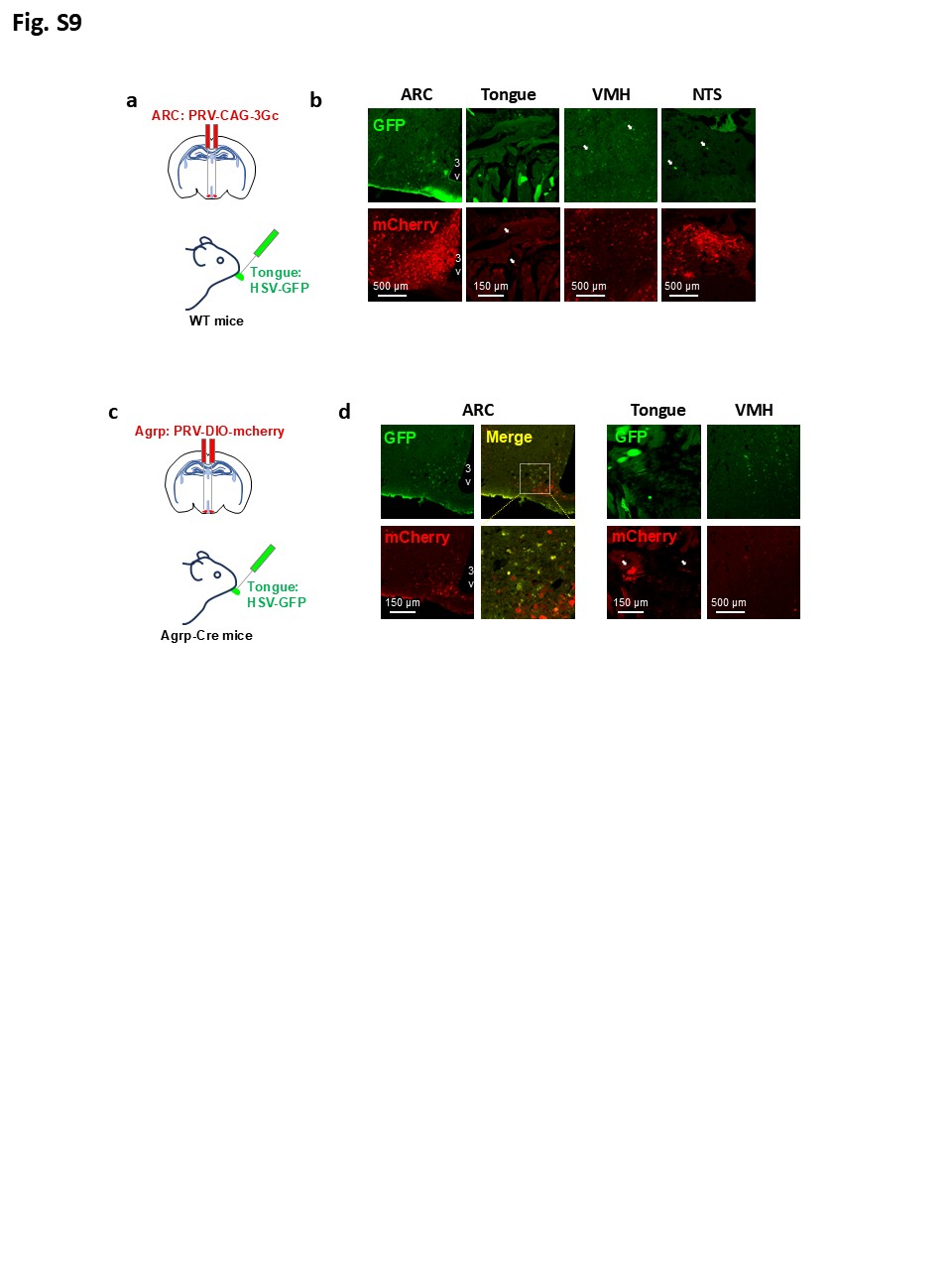

**Extended Data Fig. 9 Forward and retrograde tracing between the tongue and AgRP neurons**

a and c, Schematics illustrating the strategy of forward and retrograde tracing using HSV and PRV in wild-type (WT) or AgRP-Cre mice.

b, Immunofluorescence (IF) staining for mCherry (red), GFP (green) and Merge (yellow) in arcuate nucleus (ARC), the tongue, ventromedial nucleus (VMH) and nucleus of tractus solitarius (NTS); 3V, third ventricle.

d, IF staining for mCherry (red), GFP (green) and Merge (yellow) in ARC, the tongue, VMH.

Studies for a and b were conducted using 10-12-week-old male wild-type (WT) mice receiving forward tracing HSV (HSV-EGFP) in tongue and retrograde tracing PRV (PRV-GAG-mCherry) in ARC. Studies for c and d were conducted using 12-week-old male AgRP-Cre mice receiving forward tracing HSV (HSV-EGFP) in tongue and Cre-dependent retrograde-tracing PRV (PRV-DIO-GAG-mCherry) in ARC.

**Supplementary Table 1. Primers used for gene amplification.**

| **Gene** | **Direction** | **Primer sequence 5’→3’** |
| --- | --- | --- |
| *Gad1* | F  R | CACAGGTCACCCTCGATTTTT  ACCATCCAACGATCTCTCTCATC |
| *Slc32a1* | F  R | ACCTCCGTGTCCAACAAGTC  CAAAGTCGAGATCGTCGCAGT |
| *Npy* | F  R | CTCGTGTGTTTGGGCATT C  GATTGATGTAGTGTCGCAGAG |
| *Npy1r* | F  R | TGATCTCCACCTGCGTCAAC  ATGGCTATGGTCTCGTAGTCAT |
| *Npy2r* | F  R | GCCAGGGCACACTACTCCTA  CTACCCCTAGCAAGATGATGGA |
| *Npy5r* | F  R | TTTGTCACGGAGAACAATACTGC  TGCGCTTTTTCATAACAGCCAT |
| *Gabra2* | F  R | GGACCCAGTCAGGTTGGTG  TCCTGGTCTAAGCCGATTATCAT |
| *Gabrb3* | F  R | CTGCTGCCAATCTGGCTTTC  CGTAGCCTTTCAACAGCTTGTC |
| *Gabbr1* | F  R | TGGTTTCTCATCGGGTGGTAT  CCAAGGCCCAGATAGCATCA |
| *Gabbr2* | F  R | AAGACCCCATAGAGGACATCAA  GGGTGGTACGTGTCTGTGG |
| *Mc3r* | F  R | TCCGATGCTGCCTAACCTCT  GGATGTTTTCCATCAGACTGACG |
| *Mc4r* | F  R | ATTTGCAGCCTGCTTTCCA  TGGAGCGCGTAAAAGATAGTGAA |
| *Agrp* | F  R | TAGATCCACAGAACCGCGAGT  GAAGCGGCAGTAGCACGTA |
| *Gcn2* | F  R | CCTGCACCATGAGAACATTG  CTGCCCAGTTCTTCAGTGT |
| *Asns* | F  R | CCGTCAGATCTTTGAACGCC  TGAGTCAGCCAATCAGCCC |
| *LeuWAG* | F  R | GGTAGYGTGGCCGAGCG  GAGAATTCCATGGCAGYGGTGGG |
| *Gapdh* | F  R | TGTGTCCGTCGTGGATCTGA  CCTGCTTCACCACCTTCTTGAT |
